# Comparing RADseq and microsatellites to infer complex phylogeographic patterns, a real data informed perspective in the Crucian carp, *Carassius carassius*, L

**DOI:** 10.1101/025973

**Authors:** Daniel L Jeffries, Gordon H Copp, Lori Lawson Handley, K. Håkan Olsén, Carl D Sayer, Bernd Hänfling

## Abstract

The conservation of threatened species must be underpinned by phylogeographic knowledge in order to be effective. This need is epitomised by the freshwater fish *Carassius carassius*, which has recently undergone drastic declines across much of its European range. Restriction Site Associated DNA sequencing (RADseq) is being increasingly used for such phylogeographic questions, however RADseq is expensive, and limitations on sample number must be weighed against the benefit of large numbers of markers. Such tradeoffs have predominantly been addressed using simulated data. Here we compare the results generated from microsatellites and RADseq to the phylogeography of *C. carassius*, to add real-data-informed perspectives to this important debate. These datasets, along with data from the mitochondrial cytochrome b gene, agree on broad phylogeographic patterns; showing the existence of two previously unidentified *C. carassius* lineages in Europe. These lineages have been isolated for approximately 2.2-2.3 M years, and should arguably be considered as separate conservation units. RADseq recovered finer population structure and stronger patterns of IBD than microsatellites, despite including only 17.6% of samples (38% of populations and 52% of samples per population). RADseq was also used along with Approximate Bayesian Computation to show that the postglacial colonisation routes of *C. carassius* differ from the general patterns of freshwater fish in Europe, likely as a result of their distinctive ecology.

## Introduction

Phylogeographic studies have revealed that the contemporary distributions of European taxa and their genetic diversity have been largely shaped by the glacial cycles of the Pleistocene epoch, and in particular by range shifts during recolonisation from glacial refugia(Hewitt 1999). In freshwater fishes, the dynamics of recolonisation are tightly linked to the history of river drainage systems(Bianco 1990; Bănărescu 1990, 1992; Bernatchez & Wilson 1998; Reyjol *et al*. 2006). For example, watersheds pose a significant barrier to fish dispersal, often resulting in strong genetic structuring across contiguous drainage systems but during glacial melt periods, ephemeral rivers and periglacial lakes can arise, providing opportunities for colonisation(Gibbard *et al*. 1988) of otherwise isolated drain basins(Grosswald 1980; Arkhipov *et al*. 1995). These processes have resulted in complicated recolonisation scenarios in Europe, which, in contrast to North America(Bernatchez & Wilson 1998), appear to possess few general patterns of population structure in European fishes(Costedoat & Gilles 2009). The lack of obvious European pattern could be explained, at least in part, by the focus of phylogeographic studies on highly mobile, obligatory or facultative lotic species, with more sedentary, lentic species being largely overlooked.

The crucian carp, *Carassius carassius* (Linnaeus 1758), is native to parts of central, eastern and northern Europe and almost exclusively restricted to lentic ecosystems, including lakes, ponds and river floodplains (Copp 1991; Copp *et al*. 2008). *C. carassius*, has recently experienced sharp declines in the number and sizes of populations throughout its native range, leading to some local population extinctions. The reasons for these declines include habitat loss through drought and terrestrialisation in England (Copp 1991; Wheeler 2000; Sayer *et al*. 2011), acidification (Holopainen & Oikari 1992), poor water quality in the Danube river catchment (Navodaru *et al*. 2002), and hybridisation with several non-native species (Copp *et al*. 2010; Savini *et al*. 2010; Mezhzherin *et al*. 2012; Wouters *et al*. 2012; Rylková *et al*. 2013). The susceptibility of *C. carrassius* to genetic isolation and bottlenecks is compounded by small population sizes (Hänfling *et al*. 2005) and low dispersal (Holopainen *et al*.). Strong geographic structure is therefore likely in this species. Although the threats to *C. carassius* populations are recognised on a regional level (Lusk *et al*. 2004; Mrakovčić *et al*. 2007; Wolfram & Mikschi 2007; Simic, V *et al*. 2009; Copp & Sayer 2010), a global conservation strategy is missing. Broad scale phylogeographic data and definition of evolutionary significant units are essential for informing unified conservation efforts for this species.

Phylogeographic data have traditionally been collected using mitochondrial gene regions and/or nuclear markers such as AFLPs and microsatellites. However, cost and time often limits the number of these nuclear markers used, which can result in low power for addressing phylogeographic questions (Cornuet & Luikart 1996; Luikart & Cornuet 2008; Landguth *et al*. 2012; Peery *et al*. 2012; Hoban *et al*. 2013). Single nucleotide polymorphisms (SNPs) are increasingly used in phylogeography for assessments of population structure (for example see Morin *et al*. 2010; Emerson *et al*. 2010; Hess *et al*. 2011; Hauser *et al*. 2011) which provide several advantages (Morin *et al*. 2004). One disadvantage of this approach, however, is that bi-allelic SNP loci contain less information than the highly polymorphic AFLPs or microsatellites (Coates *et al*. 2009). Large numbers of SNPs are consequently needed to provide adequate statistical power. SNP discovery and assay development, which has been costly and slow in the past, has recently been greatly facilitated by Restriction Site-Associated DNA sequencing (RADseq, (Miller *et al*. 2006)), which enables thousands of orthologous SNP markers to be quickly isolated from non-model organisms. Despite this new opportunity, microsatellites may still be still more informative and/or cost effective in many cases, allowing for wider geographic coverage and sampling of more individuals per population. A comparison of the utility of RADseq-derived SNPs and microsatellites for phylogeographic studies is needed and will contribute to the important debate on whether it is more advantageous to genotype small numbers of highly polymorphic markers in a large number of samples, or tens of thousands of SNP markers in fewer samples. This trade-off has recently been highlighted as among the most important questions in landscape genetics (Epperson *et al*. 2010; Balkenhol & Landguth 2011).

The optimal phylogeographic study design depends heavily on the properties of the study system; in particular the strength of population structure (i.e. *F*_ST_). In *C. carassius* we expect population structure to be strong and driven by isolation by distance (IBD). If this is so, then patchy geographic sampling along the IBD gradient could result in falsely identified distinct lineages (Schwartz & McKelvey 2009). We would, therefore, expect the number of populations sampled and their geographic uniformity to be more important than number of loci, or number of samples per population in this study.

In the present study, we use a combination of mitochondrial DNA (mtDNA), microsatellites and genome-wide single nucleotide polymorphisms (SNPs) obtained from RADseq in order to: 1) produce a range wide phylogeography for *C. carassius* as a basis for Europe-wide conservation strategies, 2) test competing scenarios that have potentially contributed to the contemporary distribution of the species, and 3) compare the power of microsatellites and RADseq based population structure analyses, in the context of the first two objectives. In this third aim, we add perspectives from real biological data to a topic that has almost exclusively been addressed using simulated datasets (but see (Coates *et al*. 2009; Hess *et al*. 2011). Specifically we ask, whether the benefits gained by the high numbers of markers obtained from RADseq outweigh the potential loss of power associated by the reduction in the number of samples.

## Methods

### Sample collection and DNA extraction

We collected 848 *C. carassius* tissue samples from 49 populations across the species’ distribution in central and northern Europe (Table 1, Figure 1). Sample sizes ranged from n=4 to n=37, with a mean of n=17 (Table 1). Fish were anaesthetised by a UK Home Office (UKHO) personal license holder (GHC) in a 1 mL L^-1^ bath of 2-phenoxyethanol prior to collection of a 1 cm^2^ tissue sample from the lower-caudal fin, and wounds treated with a mixture of adhesive powder (Orahesive) and antibiotic (Cicatrin)(Moore *et al*. 1990). Tissue samples were immediately placed in ≥95% ethanol, and stored at -20^o^C. DNA was extracted from 2–4 mm^2^ of each tissue sample using either the Gentra Puregene DNA isolation kit or the DNeasy DNA purification kit (both Qiagen, Hilden, Germany). For the RADseq library, DNA was quantified using the Quant-iT™ PicoGreen® dsDNA Assay kit (Invitrogen) and normalised to concentrations ≥50 ng ml^-1^. Gel electrophoresis was then used to check that DNA extractions contained high molecular weight DNA.

**Table 1.** Location, number, genetic marker sampled, and accession numbers of samples and sequences used in the present study for microsatellite and mitochondrial DNA analyses. mtDNA sequence accession numbers can be found in Supplementary Table 5.

| Code | Location | Country | Drainage | Coordinates |  | Microsatellites<br>(n) | mtDNA<br>(n) | RAD-seq<br>(n) |
| --- | --- | --- | --- | --- | --- | --- | --- | --- |
|  |  |  |  | lat | long |  |  |  |
| GBR1 | London | U.K. | U.K | 51.5 | 0.13 | 9 |  |  |
| GBR2 | Reading | U.K. | U.K | 51.45 | -0.97 | 4 |  |  |
| GBR3 | Norfolk | U.K. | U.K | 52.86 | 1.16 | 7 |  |  |
| GBR4 | Norfolk | U.K. | U.K | 52.77 | 0.75 | 27 |  | 9 |
| GBR5 | Norfolk | U.K. | U.K | 52.77 | 0.76 | 14 |  |  |
| GBR6 | Norfolk | U.K. | U.K | 52.54 | 0.93 | 29 | 3 |  |
| GBR7 | Norfolk | U.K. | U.K | 52.9 | 1.15 | 24 | 1 | 10 |
| GBR8 | Hertfordshire | U.K. | U.K | 52.89 | 1.1 | 37 | 3 | 9 |
| GBR9 | Norfolk | U.K. | U.K | 52.8 | 1.1 | 27 |  |  |
| GBR10 | Norfolk | U.K. | U.K | 52.89 | 1.1 | 14 |  |  |
| GBR11 | Norfolk | U.K. | U.K | 52.92 | 1.16 | 20 |  |  |
| BEL1 | Bokrijk | Belgium | Scheldt River | 50.95 | 5.41 | 13 | 1 |  |
| BEL2 | Meer van Weerde | Belgium | Scheldt River | 50.97 | 4.48 | 12 |  |  |
| BEL3 | Meer van Weerde | Belgium | Scheldt River | 50.97 | 4.48 | 8 |  |  |
| GER1* | Kruegersee | Germany | Elbe River | 52.03 | 11.97 |  | 3 |  |
| GER2 | Münster | Germany | Rhine River | 51.89 | 7.56 | 21 | 3 |  |
| GER3 | Bergheim | Germany | Danube River | 48.73 | 11.03 | 9 | 3 |  |
| GER4 | Bergheim | Germany | Danube River | 48.73 | 11.03 | 8 | 3 |  |
| CZE1 | Lužnice | Czech Republic | Danube River | 48.88 | 14.89 | 9 | 3 |  |
| POL1 | Sarnowo | Poland | Vistula River | 52.93 | 19.36 | 33 |  |  |
| POL2 | Kikót-Wies | Poland | Vistula River | 52.9 | 19.12 | 34 |  |  |
| POL3 | Tupadly | Poland | Vistula River | 52.74 | 19.3 | 17 |  | 10 |
| POL4 | Orzysz | Poland | Vistula River | 53.83 | 22.02 | 13 | 3 | 10 |
| EST1 | Tartu | Estonia | Baltic Sea | 58.39 | 26.72 | 5 | 3 |  |
| EST2 | Vehendi | Estonia | Baltic Sea | 58.39 | 26.72 | 5 |  |  |
| RUS4* | Velikaya river | Russia | Baltic Sea | 55.9 | 30.25 | 29 | 3 |  |
| FIN1 | Joensuu | Finland | Baltic Sea | 62.68 | 29.68 | 32 | 3 |  |
| FIN2 | Helsinki | Finland | Baltic Sea | 60.36 | 25.33 | 32 |  |  |
| FIN3 | Jyväskylä | Finland | Baltic Sea | 62.26 | 25.76 | 37 | 3 | 10 |
| FIN4 | Oulu | Finland | Baltic Sea | 65.01 | 25.47 | 7 | 3 | 8 |
| FIN5 | Salo | Finland | Baltic Sea | 60.37 | 23.1 | 10 | 3 |  |
| SWE1 | Gränbrydampen | Sweden | Baltic Sea | 59.87 | 17.67 | 25 |  |  |
| SWE2 | Stordammen | Sweden | Baltic Sea | 59.8 | 17.71 | 21 | 3 | 10 |
| SWE3 | Östhammar | Sweden | Baltic Sea | 60.26 | 18.38 | 27 | 3 |  |
| SWE4 | Umeå | Sweden | Baltic Sea | 63.71 | 20.41 | 9 | 3 |  |
| SWE5 | Kvicksund | Sweden | Baltic Sea | 59.45 | 16.32 | 9 |  |  |
| SWE6 | Åland Island | Sweden | Baltic Sea | 60.36 | 19.85 | 8 | 3 |  |
| SWE7 | Grillby | Sweden | Baltic Sea | 59.64 | 17.37 | 10 |  |  |
| SWE8 | Skabersjö | Sweden | Baltic Sea | 55.55 | 13.15 | 19 | 3 | 10 |
| SWE9 | Märsta | Sweden | Baltic Sea | 59.6 | 17.8 | 31 | 3 |  |
| SWE10 | Norrköping | Sweden | Baltic Sea | 58.56 | 16.27 | 29 |  | 9 |
| SWE11 | Gotland Island | Sweden | Baltic Sea | 57.85 | 18.79 | 11 | 3 |  |
| NOR1 | Oslo | Norway | North Sea | 60.05 | 9.94 |  | 2 |  |
| NOR2 | Tromsø | Norway | North Sea | 69.65 | 18.95 | 16 |  | 9 |
| BLS | - | Belarus | Dnieper | 52.47 | 30.52 | 7 | 1 |  |
| RUS1 | Proran Lake | Russia | Don River | 47.46 | 40.47 | 10 | 3 | 9 |
| DEN1 | Copenhagen | Denmark | Baltic Sea | 60.21 | 17.79 | 12 |  | 10 |
| DEN2 | Pederstrup | Denmark | Baltic Sea | 55.77 | 12.55 | 14 |  | 8 |
| DEN3 | Gammel Holte | Denmark | Baltic Sea | 56 | 12.5 | 14 |  |  |
| DEN4 | Bornholm Island | Denmark | Baltic Sea | 55.17 | 14.86 |  |  | 5 |
| SWE12 | Osterbybruk | Sweden | Baltic Sea | 55.73 | 12.34 | 14 |  | 9 |
| SWE14 | Stockholm | Sweden | Baltic Sea | 59.66 | 18.95 | 16 |  | 9 |
| RUS2* | Karma | Russia | Volga River | 52.9 | 58.4 |  | 2 |  |
| RUS3* | Saygach'yedake | Russia | Volga River | 47.5 | 48.5 |  | 4 |  |
| HUN1 | Gödöllő | Hungary | Danube River | 47.61 | 19.36 |  | 2 |  |
| HUN2 | Vörösmocsár | Hungary | Danube River | 46.49 | 19.17 |  |  | 6 |
| **Total** |  |  |  |  |  | **848** | **79** | **154** |

(Table 1 continued)
| **Genbank mtDNA sequences** |  |  |  |  |
| --- | --- | --- | --- | --- |
| **Code** | **Reference** | **Country** | **Drainage** | **Accession** |
| GER6 | Kalous et al. (2007) | Germany | Baltic sea | DQ399917 |
| GER6 | Kalous et al. (2007) | Germany | Baltic sea | DQ399918 |
| GER6 | Kalous et al. (2007) | Germany | Baltic sea | DQ399919 |
| GER7 | Rylková et al. (2013) | Germany | Hunte River | JN412540 |
| GER7 | Rylková et al. (2013) | Germany | Hunte River | JN412541 |
| GER7 | Rylková et al. (2013) | Germany | Hunte River | JN412542 |
| GER7 | Rylková et al. (2013) | Germany | Hunte River | JN412543 |
| GER8* | Rylková et al. (2013) | Germany | Lahn River | JN412537 |
| GER8* | Rylková et al. (2013) | Germany | Lahn River | JN412538 |
| CZE2 | Rylková et al. (2013) | Czech Republic | Elbe drainage | GU991399 |
| AUS1 | Rylková et al. (2013) | Austria | Danube river | JN412533 |
| AUS1 | Rylková et al. (2013) | Austria | Danube river | JN412534 |
| AUS2 | Rylková et al. (2013) | Austria | Danube river | JN412535 |
| AUS3 | Rylková et al. (2013) | Austria | Danube river | JN412536 |
| GBR12 | Rylková et al. (2013) | U.K. | U.K | JN412539 |
| GBR12 | Kalous et al. (2012) | U.K. | U.K | GU991400 |
| SWE15 | Rylková et al. (2013) | Sweden | Baltic sea | JN412545 |
| SWE16 | Rylková et al. (2013) | Sweden | Baltic sea | JN412544 |

† Also present
\* Location on Map (Fig. 1.a) is approximate

**Figure 1.**
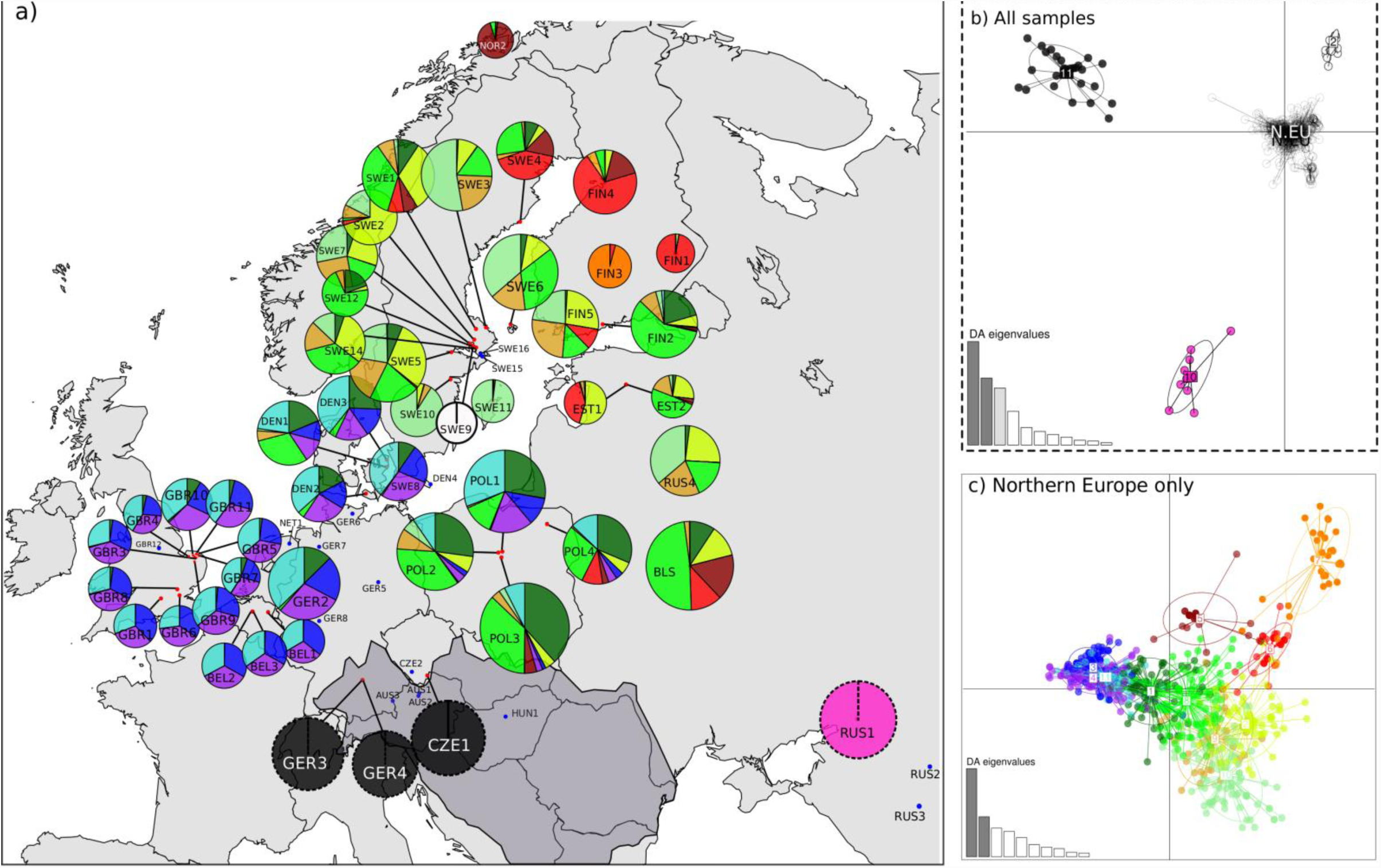
Population structure of C. carassius in Europe. a) Sampling locations (sites sampled with nuclear and mtDNA markers = red dots, mtDNA only = blue dots) and population cluster memberships from DAPC analysis. Pie chart size corresponds to microsatellite allelic richness. Pie chart colours for Danubian populations and RUS1 correspond to clusters in the broad scale DAPC analysis b) and for all northern European populations colours correspond to clusters in the northern European DAPC analysis (mtDNA lineage 1 only) c). The Danube river catchment is shaded dark grey.

### Molecular markers and methods

Three types of molecular markers were used in the study. Mitochondrial DNA sequencing was used to identify highly distinct lineages and to date the divergence between them through phylogenetic analysis. Two sets of nuclear markers; microsatellites and RADseq-derived SNPs were used to investigate more recent and complex structure in a population genetics framework and to compare the relative power of each marker to do so.

### Mitochondrial DNA amplification

A total of 82 *C. carassius* individuals, randomly chosen from a subset of 30 populations, which were chosen to represent all major catchment areas and the widest possible geographic range (min. n = 1, max. n = 4, mean n = 2.7), were sequenced at the cytochrome b (*cyt*b) gene (Table 1). PCR reactions were carried out following the protocol in Takada *et al*. (2010) *using the forward and reverse primers L14736-Glu and H15923-Thru on an Applied Biosciences® Veriti Thermal Cycler. PCR products were sequenced in both directions on an ABI3700 by Macrogen Europe. The forward and reverse cyt*b sequence reads were aligned using a GenBank sequence from the UK (accession no. JN412539, Table 1) as a reference and ambiguous nucleotides were manually edited using CodonCode aligner v.2.0.6 (CodonCode Corporation).

### Microsatellite amplification

All 848 *C. carassius* samples were genotyped at 13 microsatellite loci, which were originally designed for use in *Carassius auratus*, or *Cyprinus carpio* and cross amplify in *C. carassius* (Supplementary table 1). Six of these loci were chosen for their species diagnostic properties, allowing us to ensure that all samples used in the present study were *C. carassius* and not one of the closely-related introduced species (*C. carpio*, *C. auratus*, or *Carassius gibelio*) or their hybrids (see Supplementary text for full details of species identification and hybrid detection). Microsatellites were amplified in three multiplex PCR reactions, using the Qiagen multiplex PCR mix with manufacturer’s recommended reagent concentrations, including Q solution and 1 μl of template DNA. Primer concentrations for each locus are provided in Supplementary Table 1 and PCRs were performed on an Applied Biosciences® Veriti Thermal Cycler. The annealing temperature used was 54°C for all reactions, and all other PCR cycling parameters were set to Qiagen multiplex kit recommended values. PCR products were run on a Beckman Coulter CEQ 8000 genome analyser using a 400 bp size standard and microsatellite alleles scored using the Beckman Coulter CEQ8000 software.

### RADseq

A total of 149 individuals (16 populations, min. *n* = 8, max. *n* = 10, mean *n* = 8.9), identified as pure *C. carassius* with the diagnostic microsatellites, were used in the RADseq (Table 1). These samples were chosen to represent a wide geographic range and all major phylogeographic clusters identified using the microsatellite data. These samples were split across 13 libraries prepared at Edinburgh Genomics (University of Edinburgh, UK) according to the protocol in Davey *et al.* (2012) using the enzyme *Sbf*1. Libraries were then sequenced across five lanes of two Illumina HiSeq 2000 flowcells (Edinburgh Genomics).

## Data analyses

### Phylogenetic analysis of mtDNA

In addition to the 82 sequenced samples, we retrieved 18 published *C. carassius cyt*b sequences from GenBank which were validated through cross checking with their original publications (Table 1). Sequence alignment was performed in MEGA6 (Tamura *et al*. 2013) using default settings, and DNAsp v.5.0 (Librado & Rozas 2009) was used to calculate sequence divergence and to identify haplotypes.

Haplotypes were exported to BEAST v.1.7.5 (Drummond *et al*. 2012) for phylogenetic analyses in order to identify the major phylogenetic lineages within European *C. carassius*. The splits between the major phylogenetic clades were then dated using a relaxed molecular clock method in BEAST. The widely-used Dowling *et al*. (2002) *cyprinid cyt*b divergence rate of 1.05% pairwise sequence divergence / MY was used after converting to a per lineage value of 0.0053 mutations/site/MY for use in BEAST. Initial analyses using the GTR (Tavaré 1986) substitution model yielded multiple parameters with low estimated summary statistic (ESS) values (<100), therefore the less complex HKY (Hasegawa *et al*. 1985) substitution model, which had ESS values >200 for all parameters, was used. We used a ‘coalescent: constant size’ tree prior, which assumes an unknown but constant population size backwards in time, as recommended for intraspecific phylogenies (*BEAST Tutorial - Tree priors and dating*). MCMC chain lengths were 1 × 10^7^ with samples taken every 1000 iterations. A gamma site heterogeneity model was used, with the default of four categories. Substitution rates, rate heterogeneity and base frequencies were unlinked between each codon position to allow substitution rate to vary between them. Default values were used for all other parameters and priors.

### Population structure and diversity analyses using microsatellites

Allele dropout and null alleles in the microsatellite data were tested using Microchecker (Van Oosterhout *et al*. 2004). FSTAT v. 2.9.3.2 (Goudet 2001) was then used to check for linkage disequilibrium (LD) between loci (using 10,000 permutations), deviations from Hardy-Weinberg equilibrium (HWE) within populations (126500 permutations) and for all population genetic summary statistics. Genetic diversity within populations was estimated using Nei’s estimator of gene diversity (*H*_o_) (Nei 1987) and Allelic richness (*A*_r_), which was standardised to the smallest sample size (n =5) using rarefaction (Petit *et al*. 1998). Pairwise *F*_ST_ values were calculated according to (Weir & Cockerham 1984) and 23520 permutations and sequential Bonferroni correction were used to test for significance of *F*_ST_.

IBD was investigated using a Mantel test in the adegenet v1.6(Jombart & Ahmed 2011) package in R v3.0.1(R Core Team 2013). We then tested for an association between *A*_*r*_ and longitude and latitude, which is predicted under a stepping-stone colonisation model (Ramachandran *et al*. 2005; Simon *et al*. 2014), using linear regression analysis in R.

Population structure was then further examined using Discriminant Analyses of Principal Components (DAPC) also in adegenet (DAPC, see Supplementary text and Jombart *et al*. 2010 for more details). In preliminary DAPC analysis using all 49 *C. carassius* populations, Sweden (SWE9) was found to be so genetically distinct from the rest of the data set that it masked the variation between the other populations. This population was therefore omitted from further DAPC analyses. To infer the appropriate number of genetic clusters in the data was, we used Bayesian Information Criteria (BIC) scores, in all cases choosing lowest number of genetic clusters from the range suggested. Spline interpolation(Hazewinkel 1994) was then used to identify the appropriate number of principal components to use in the subsequent discriminant analysis.

### RADseq data filtering and population structure analysis

The quality of the RADseq raw read data was examined using FastQC(Andrews 2010), the dataset was then cleaned, processed and SNPs were called using the Stacks pipeline(Catchen *et al*. 2013). One SNP per RAD locus was used and, SNPs were only retained if they were present in at least 17 out of the 19 populations in the study, which allows for mutations in restriction sites that may cause loci to dropout in certain lineages. Finally, we filtered out loci which had a heterozygosity of > 0.5 in one or more populations in order to control for the possibility of erroneously merging ohnologs resulting from the multiple genome duplications that have occurred the *Cyprinus* and *Carassius* genera (Henkel *et al*. 2012; Xu *et al*. 2014). The resulting refined SNP set was then used in subsequent phylogeographic analyses. The adegenet R package was used to calculate *H*_o_ and pairwise *F*_ST_, test for IBD and infer genetic clusters using DAPC.

### Reconstructing postglacial colonisation routes in Europe

DIYABC (Cornuet *et al*. 2014) was used to reconstruct the most likely *C. carassius* recolonisation routes through Europe after the last glacial maximum. Analyses were performed on1000 randomly-selected SNP loci from the full RAD-seq dataset were used, as microsatellite loci are likely to be affected by homoplasy over the time scales used here (Morin *et al*. 2004). The reduced dataset was first analysed with DAPC to confirm that it produced the same structure as the full dataset. Then datasets of expected summary statistics were simulated for a number of scenarios (i.e. a specific population tree topology, together with the parameter prior distributions that are associated with it). These simulated datasets represent the theoretical expectation under each scenario, and are compared to the same summary statistics calculated from the observed data to identify the most likely of the specified scenarios. In DIYABC, two methods of comparison between simulated and observed datasets are used; logistic regression and “direct approach”, the latter method identifies the scenario that produces the largest proportion of the *n* number of closest scenarios to the observed, where *n* is specified by the user. The goodness-of-fit of scenarios was also assessed using the model checking function implemented in DIYABC (Cornuet *et al*. 2014).

To reduce the number and complexity of possible scenarios, we split DIYABC analysis into three stages (Table 2). In stage 1, we tested 11 broad scale scenarios (Scenarios 1 -11, Supplementary Figure 1), in which populations were grouped into three pools; Pool 1 – all northern European populations (npops = 17, *n* = 155), Pool 2 – Don population (npops = 1, *n* = 9), Pool 3 – Danubian population (npops = 1, *n* = 6). Both population pooling and scenarios were chosen on the basis of the broad phylogeographic structure identified in the mtDNA and RAD-seq population structure analysis (see Results). We tested the likelihood of these 11 scenarios, simulating one million summary-statistic datasets per scenario, for comparison to the real dataset.

**Table 2.** Population pools, parameter priors used and posterior parameter values inferred in the three stages of DIYABC analysis.

| Analysis stage | Population Pools | Scenarios tested | Parameter priors | Most likely Scenario | Median of posterior distributions of most likely scenario |
| --- | --- | --- | --- | --- | --- |
| 1 | **Pool 1** – GBR4, GBR7, GBR8, DEN1, DEN2, DEN3, FIN3, FIN4, POL3, POL4, SWE2, SWE8, SWE9, SWE10, SWE12, SWE14, NOR2 | 1 – 11 | N1 = 10E+03 - 500E+03<br>Nb1 = 10 - 100E+03<br>N2 = 100 - 100E+03<br>N3 = 100 - 200E+03<br>t1 = 1E+03 - 1E+06 gens<br>t2 = 1E+03 - 3E+06 gens<br>ra = 0.001-0.999<br>rb = 0.001-0.999<br>rc = 0.001-0.999<br>db = 10- 10E+03 gens | 9 | N1 = 3.47E+04<br>Nb1 = 2.37E+04<br>N2 = 7.49E+04<br>N3 = 1.40E+05<br>t1 = 1.35E+05<br>db = 4.46E+03<br>t2 = 1.09E+06 |
|  | **Pool 2** – RUS1 |  |  |  |  |
|  | **Pool 3** – HUN2 |  |  |  |  |
| 2 | **Pool 1** – GBR4, GBR7, GBR8 | 12 – 16 | N1 = 10-4E+03<br>N2 = 10 - 10E+03<br>N3 = 10 - 20E+03<br>N4 = 10 - 50E+03<br>N5 = 10 - 20E+03<br>N6 =10 - 400<br>t1 = 100- 10E+03 gens<br>t1a = 100- 10E+03 gens<br>t2 =100- 10E+03<br>t2a =100- 5E+03 gens<br>t2b = 500-20E+03 gens<br>t2c = 100 - 10E+03 gens<br>t2d = 100 - 10E+03 gens<br>t3 = 500 - 20E+03 gens<br>t3c =100 - 10E+03 gens<br>t3d =100 - 10E+03 gens<br>t4 =500 - 20E+03 gens<br>ra = 0.001-0.999<br>rb = 0.001-0.999 | 14 | N1 = 3.67E+03<br>N2 = 7.52E+03<br>N3 = 1.74E+04<br>N4 = 1.94E+04<br>N5 = 1.18E+04<br>N6 = 2.10E+02<br>t1 = 6.79E+03<br>t1a = 2.51E+03<br>t2d = 6.78E+03<br>t3d = 8.91E+03<br>t4 = 1.20E+04<br>rb = 6.68E-01 |
|  | **Pool 2** – DEN1, DEN2, DEN3 |  |  |  |  |
|  | **Pool 3** – FIN3, FIN4 |  |  |  |  |
| 3 | **Pool 4** – POL3, POL4 | 14a- 14f | N1 = 10-4E+03<br>Nb1 = 10-10E+03<br>N2 = 10 - 10E+03<br>N3 = 10 - 20E+03<br>Nb3 = 10-10E+03<br>N4 = 10 - 50E+03<br>N5 = 10 - 20E+03<br>N6 =10 - 400<br>Nb6 =10-10E+03<br>t1 = 100- 10E+03 gens<br>t1a = 100- 10E+03 gens<br>t2d = 100 - 10E+03 gens<br>t3d = 100 - 10E+03 gens<br>t4 = 500 - 20E+03 gens<br>rb = 0.001-0.999<br>da = 10 - 10E+03 gens<br>db = 10 - 10E+03 gens<br>dc = 10 - 10E+03 gens<br>dd = 10 - 10E+03 gens<br>de = 10 - 10E+03 gens | 14d | N1 = 2.39E+03<br>Nb1 = 9.35E+02<br>N2 = 8.14E+03<br>N3 = 9.36E+03<br>N4 = 1.70E+04<br>N5 = 1.10E+04<br>N6 = 1.38E+02<br>t1 = 3.75E+03<br>t1a = 2.46E+03<br>t2d = 5.90E+03<br>t3d = 7.97E+03<br>t4 = 1.68E+04<br>rb = 6.19E-01<br>dc = 9.07E+03 |
|  | **Pool 5** – SWE2, SWE8, SWE9, SWE10, SWE12, SWE14 |  |  |  |  |
|  | **Pool 6** – NOR2 |  |  |  |  |

In the second and third stages, we performed a finer scale analysis, focussing on the 17 northern European populations alone. Populations were again pooled on the basis of both population structure and geography, in order to reduce scenario complexity (Table 2). In stage 2 we tested five scenarios (Scenarios 12-16. Supplementary Figure 2a), with no bottlenecks included, which represented the major topological variants that were most likely, given population structure results from DAPC. We then identified the most likely of these scenarios in DIYABC and took this forward into the final stage of the analysis where we tested 6 multiple bottleneck combinations (Supplementary Figure 2b) around this scenario. This three stage approach allowed us to systematically build a complex scenario for the European colonisation of *C. carassius*. Finally, we used the posterior distributions of the time parameters from the scenario identified as most likely in stages one and three to estimate times of the major lineage splits in European *C. carassius*.

### Comparison of microsatellite and RADseq data

Finally, we compared the results derived from population structure analyses on microsatellite and RADseq data to assess their suitability for addressing our phylogeographic question. It is important to note that differences between the full microsatellite and RADseq datasets could be attributable to one or a combination of the following; the number of populations, the geographic distribution of populations, the number of samples per population, the number of markers, or the information content of the marker type. To disentangle these sources of variation, we created two microsatellite data subsets; M2, which included only individuals used in RADseq, (excluding three individuals for which microsatellite data was incomplete, *n* = 146, npops = 19), and M3, which contained all individuals for which microsatellite data was available in populations that were used in RADseq (*n* = 313, npops = 19; Table 3). This gave us three pairs of datasets for comparison: 1) RADseq Vs. M2: same individuals but different marker types, 2) M1 vs M2: full microsatellite dataset versus a subset of the populations, and 3) M2 vs M3: same populations but different number of individuals per population. This strategy enabled us to test for the influence of marker, sampling of populations and individuals per population respectively. Comparisons were performed between datasets on heterozygosities and pairwise *F*_ST_s using both Pearson’s product-moment correlation coefficient and paired Student’s t-tests in R. IBD results were compared using Mantel tests (Jombart & Ahmed 2011), and DAPC results were compared on the basis of similarity of number of inferred clusters and cluster sharing between populations.

**Table 3.** Summary statistics for M1, M2, M3 and RADseq datasets. RAD contains all RAD-seq data, M1 contains all microsatellite data, M2 contains only microsatellite for the individuals used in the RAD-seq, and M3 contains all microsatellite data for all individuals that were available in populations that were used in RAD-seq.

| Dataset | Description | N samples | Mean N samples/pop | N. loci | Mean N.alleles/pop | Mean N.alleles/locus |
| --- | --- | --- | --- | --- | --- | --- |
| RAD | RADseq data only | 149 | $8.95 \pm 1.4$ | 13189 | 6723 | 2 |
| M1 | Full Microsatellite dataset | 848 | $17.2 \pm 9.5$ | 13 | $27 \pm 8.8$ | 7.6 |
| M2 | Microsatellites for RADseq samples only | 146 | $9.125 \pm 0.8$ | 13 | $24.4 \pm 7.3$ | $7.8.4 \pm 5.1$ |
| M3 | Microsatellites for all samples in populations used in RADseq | 313 | $19.6 \pm 9.0$ | 13 | $27.4 \pm 8.1$ | $11.23 \pm 7.6$ |

## Results

### Phylogenetic analyses of Mitochondrial data

The combined 1090 bp alignment of 100 *cyt*b *C. carassius* mtDNA sequences yielded 22 haplotypes, which were split across two well supported and highly differentiated phylogenetic lineages (Figure 2, Supplementary table 2). Lineage 1 was found in all northern European river catchments sampled, as well as eastern European (Dnieper) and southeastern European (Don and Volga) catchments, whereas Lineage 2 was almost exclusively confined to the River Danube catchment. There were a few exceptions to this clear geographical split however; two individuals, one from the Elbe and one from the Rhine in northern Germany, belonged to mtDNA Lineage 2, as did one individual from the River Lahn river catchment in western Germany. Also one population in the Czech Republic, located on the border between the Danube and Rhine river catchments, was found to contain individuals belonging to lineages 1 and 2.

**Figure 2.**
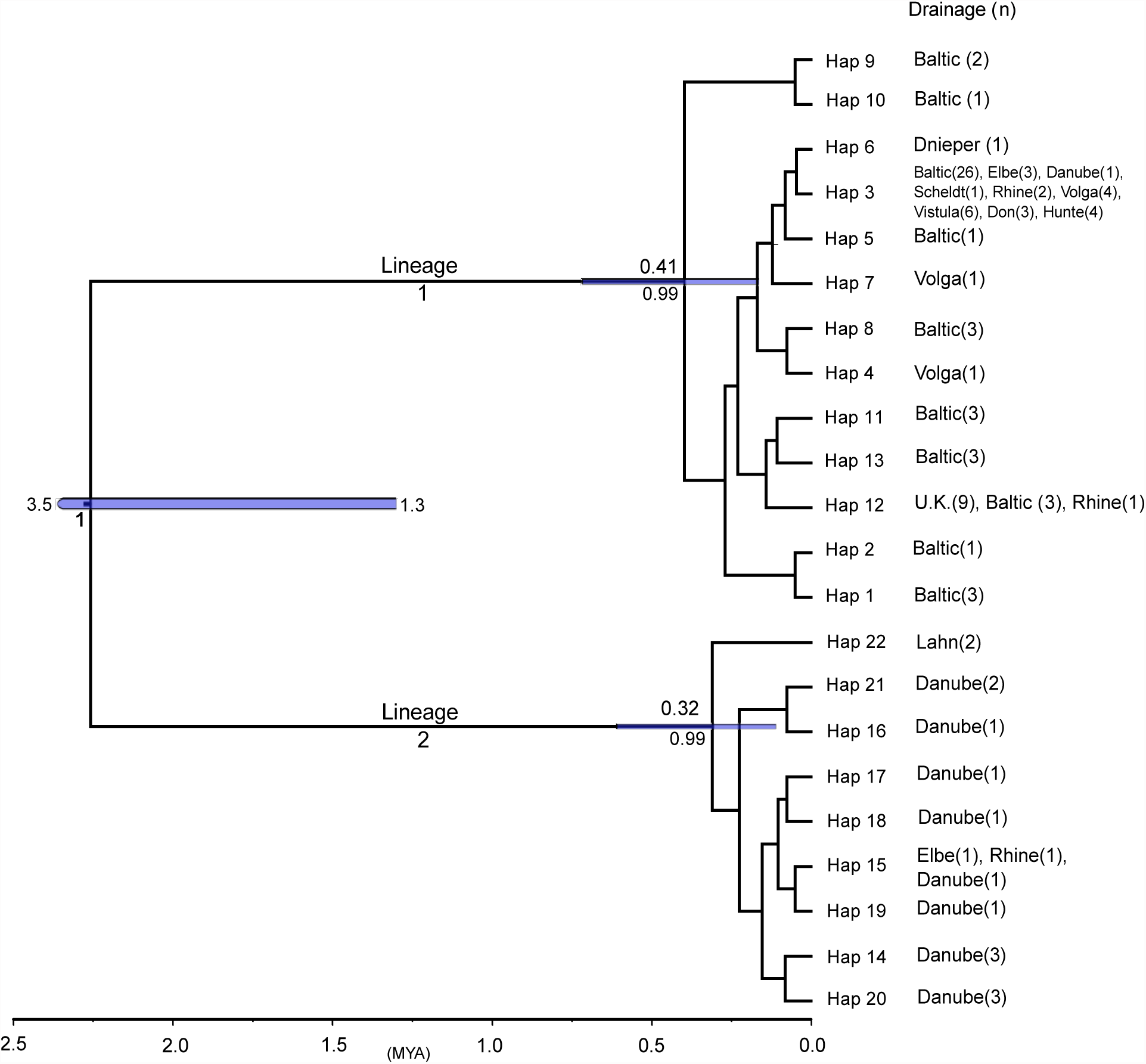
Maximum credibility tree calculated in BEAST for 100 C. carassius cytb sequences. For the three maximally supported nodes, age is given above and the posterior probability distribution is given below, with 95% CI’s represented by blue bars.

The mean number of nucleotide differences within lineages 1 and 2 was 2.25 and 2.00, respectively, which equated to a sequence divergence 0.2% and 0.18%, respectively. Between the two lineages there was an average of 22.5 nucleotide differences (2.06% mean sequence divergence), with 19 of these being fixed. BEAST molecular clock analysis dated the split between lineages 1 and 2 to be 1.30–3.22 million years ago (MYA), with a median estimate of 2.26 MYA (Figure 2).

### Nuclear marker datasets and quality checking

Microchecker showed no consistent signs of null alleles or allele dropout in microsatellite loci and no significant LD was found between any pairs of loci. No populations showed significant deviation from Hardy-Weinberg proportions (adjusted nominal level 0.0009).

After filtering raw RADseq data, *de novo* construction of loci across the 19 populations produced 35 709 RADseq loci that were present in at least 70% of individuals in at least 17 populations. These loci contained a total of 29 927 polymorphic SNPs (approx. 0.84 SNPs per locus). Only the first SNP in each RADseq locus was retained, to avoid confounding signals of LD. This yielded a total of 18 908 loci with a mean coverage of 29.07 reads. Finally 5719 of these SNP loci were filtered out due to high (> 0.5) heterozygosity in at least one population. In doing so, we removed many high coverage tags (Supplementary Fig. 3), which was consistent with over-merged ohnologs having higher coverage (i.e. reads from more than two alleles) than correctly assembled loci. The final dataset therefore contained 13189 SNP loci, with a mean coverage of 27.72 reads.

### Within population diversity at nuclear loci

Observed heterozygosity (*H*_o_), averaged across all microsatellite loci within a population, ranged from 0.06 (SWE9) to 0.44 (BLS), with a mean of 0.25 across all populations (SD = 0.105), and was highly correlated with *A*_r_ (*t* = 19.67, *P* < 0.001, df = 40), which ranged from 1.26 (FIN1) to 2.96 (POL3) with a mean of 1.92 (SD = 0.51). Mean *H*_o_ averaged across all RADseq loci for all populations was 0.013 (SD = 0.013), ranged from 0.001 to 0.057 and was significantly correlated with *H*_o_ from microsatellite loci at populations shared between both datasets (r = 0.69, *t* = 3.74, *P* = 0.002, df = 15). Microsatellite *A*_r_ significantly decreased along an east to west longitudinal gradient (adj. *R*^2^ = 0.289, *P* < 0.001, Supplementary Figure 4b) consistent with decreasing diversity along colonisation routes. However, *A*_r_ did not decrease with increasing latitude (Adj R2 =-0.007, P = 0.414, Supplementary Figure 4a). We also repeated this analysis after removing samples from mtDNA Lineage 2 in the Danube catchment. Again there was no relationship between *A*_r_ and latitude (*R*2 =-0.023, *P* = 0.254, Supplementary Figure 4c), but the relationship between *A*_r_ and longitude was strengthened (adj. *R*^2^ = 0.316, *P* < 0.001, Supplementary Figure 4d).

### Population Structure in Europe based on nuclear markers

Population structure was strong, as predicted. Using the full (M1) microsatellite dataset, mean pairwise *F*_ST_ was 0.413 (min = 0.0; BEL2 and BEL3), max = 0.864 (NOR2 vs GBR2), with 861 of the 1128 pairwise population comparisons being significant *F*_ST_ (*P* < 0.05, Supplementary table 3). Pairwise *F*_ST_ calculated from the RADseq dataset also showed strong structure (Supplementary table 4), ranging from 0.067 (DEN1, DEN2) to 0.699 (NOR2, GBR4), and these values were highly correlated with the same population comparisons in the M3 microsatellite dataset (r = 0.66, *t* = 9.01, *P* < 0.01, df = 104).

BIC scores obtained from initial DAPC analyses, using all 49 populations, indicated that between 11 and 19 genetic clusters (Supplementary Figure 5a) would be an appropriate model of the variation in the data. As a conservative estimate of population structure, we chose 11 clusters for use in the discriminant analysis, retaining eight principal components as recommended by the spline interpolation a-scores (Supplementary Figure 5a). This initial analysis showed that populations belonging to Cluster 10 (RUS1, Don river catchment) and Cluster 11 (GER3, GER4, CZE1, Danubian catchment) were highly distinct from clusters found in northern Europe (Figure 1b). Since the marked genetic differentiation between these three main clusters masked the more subtle population structure among northern European populations (see Figure 1b), we repeated the DAPC analysis without the populations from the Danube and Don (RUS1, GER3, GER4, CZE1, Figure 1b). The results of this second DAPC analysis revealed an IBD pattern of population structure, across Europe (Figure 1). Mantel tests excluding the Danubian and Don populations corroborated these results; showing significant correlation with geographic distance in northern Europe (adjusted *R*^2^ = 0.287, *P* < 0.001, Supplementary Figure 6a), with Danubian populations shown to be more diverged than their geography would predict (data not shown).

In the RADseq DAPC analysis, BIC scores suggested between four and ten genetic clusters, similar to the range inferred in the microsatellite data, and we therefore chose four clusters to take forward in the analysis (Supplementary Figure 5b). Following spline interpolation, we retained six principal components and kept two of the linear discriminants from the subsequent discriminant analysis (Supplementary Figure 5b). The inferred population structure showed that the Danubian population (HUN2) and the Don population (RUS1) were highly diverged from the northern European clusters. Unfortunately, HUN2 is not present in the microsatellite dataset for direct comparison, however both datasets, and the mtDNA data show the same pattern of high divergence between northern Europe and Danubian populations. DAPC analyses of RADseq data again showed an IBD pattern in northern European populations, which was confirmed with Mantel tests when the Danubian population HUN2 was excluded (adjusted *R*^2^ = 0.722, *P* < 0.001; Supplementary Figure 6b).

### Postglacial recolonisation of C. carassius in Europe

DAPC results of the 1000 SNP RADseq dataset used in DIYABC showed that it produced the same population structure as the full RADseq dataset (Supplementary Figure 7). For the broad-scale scenario tests in stage one of the DIYABC analysis, both logistic regression and direct approach identified Scenario 9 as being most likely to describe the true broad-scale demographic history (Supplementary Figure 8). Model checking showed that the observed summary statistics for our data fell well within those of the posterior parameter distributions for scenario 9 (Supplementary Figure 8c). Scenario 9 agrees with the mtDNA results, suggesting that the Danubian populations have made no major contribution to the colonisation of northern Europe. The median posterior distribution estimate of the divergence time between Danubian and northern European populations is 2.18 MYA (assuming a two-year generation time;(Tarkan *et al*. 2010), which is strikingly similar to that of mtDNA dating analysis. Scenario 9 also suggests that the northern European populations experienced a population size decline after the split of Pool 1 from the population in the Don river catchment, which lasted approximately 8920 years and reduced *N*_*e*_ by 32%.

In stage two of the DIYABC analysis, we tested the major variant scenarios for the colonisation of northern Europe. In assessing the relative probabilities of scenarios, there was some discrepancy between the direct approach, which revealed Scenario 14 to be most likely, and the logistic regression, which favoured Scenario 13 (with Scenario 14 being the second most likely). However, the goodness-of-fit model checking showed that the observed dataset fell well within the posterior parameter distributions for Scenario 14 (Supplementary Figure 9a), but not for Scenario 13 (not shown). Therefore, Scenario 14 was carried forward into stage three in which we tested six more scenarios (Supplementary Figure 2b) to compare combinations of bottlenecks using the same population tree topology as in Scenario 14. Direct approach, logistic regression and model checking all found scenario 14d to be the most likely (Supplementary Figure 9b), we therefore accepted this as the scenario for the colonisation of *C. carassius* in northern Europe (Supplementary Figure 9b). This scenario infers an initial split between two sub-lineages in northern Europe approximately 33 600 YBP (Figure 4), one of which re-colonised northwest Europe and one that re-colonised Finland through the Ukraine and Belarus. Scenario 14d also inferred a secondary contact between these sub-lineages approximately 15 940 YBP, resulting in the populations currently present in Poland; these admixed populations provided the source of one colonisation across the Baltic into Sweden, and a second route was inferred into southern Sweden from Denmark (Table 3, Supplementary Figure 9b).

### Comparing microsatellite datasets and RAD-sequencing data

The results from the RADseq (*n* = 149, npops = 16) dataset and the full microsatellite dataset (M1, *n* = 848, npops = 49) largely agreed on the inferred structure and cluster identity of populations. However, there were some important differences between them. Firstly, the IBD pattern of population structure in northern Europe was much stronger in the RADseq data (R^2^ = 0.722, *P* < 0.001) (Supplementary Figure 6) compared to the M1 dataset (R^2^ = 0.287, *P* < 0.001) (excluding Danubian populations and SWE9 from both datasets, Supplementary Figure 6). Secondly, clusters inferred by the RADseq DAPC analysis are much more distinct, i.e. there is much lower within-cluster, and higher between-cluster variation in the RADseq results than in the M1 dataset results (Figure 3).

**Figure 3.**
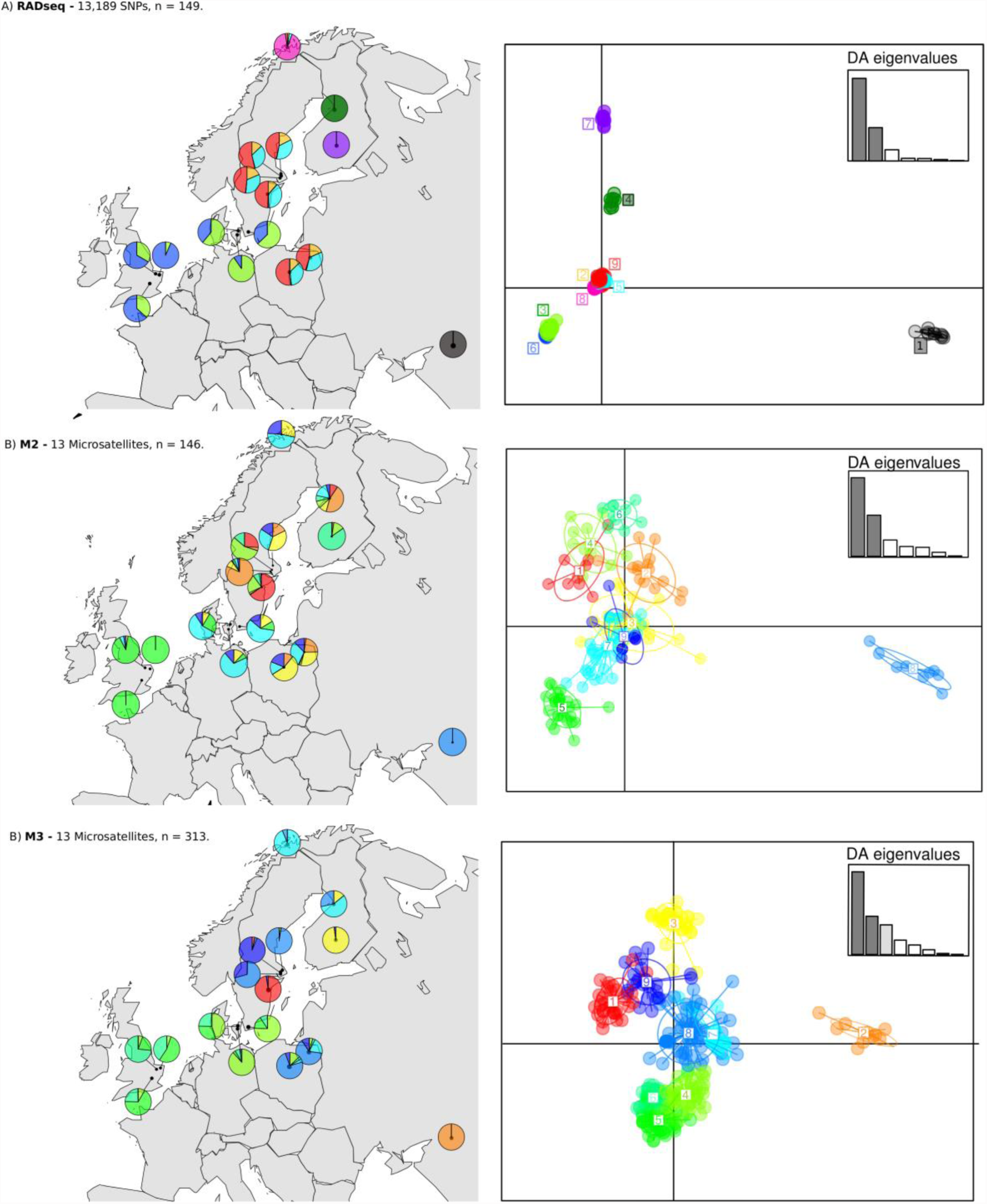
Comparison of DAPC results using RADseq dataset a), M2 dataset b) and M2 dataset c). Colours correspond between DAPC scatter plots and maps within but not between panels.

As the properties of the RADseq and M1 datasets differ in four respects, namely marker type, number of populations, number of samples per population (Table 3) and uniformity of sampling locations, (Supplementary Figure 10), it was not possible to identify the cause of discrepancies in their results. Therefore, below we report the results from the pair-wise dataset comparisons, which isolate the effects of these parameter differences.

*1) M1 Vs. M3:* the effect that the number of populations and the uniformity of sampling locations might have on inferred population structure. The geographic distribution of sampling locations was more clustered in M1 (full microsatellite dataset) than in M3 (containing microsatellite for samples in populations used in RADseq, Supplementary Figure 10), and IBD patterns were considerably stronger in the M3 subset (adj. *R*_2_ = 0.447, *P* < 0.001) than in the full M1 dataset (adj. *R*_2_ = 0.287, *P* < 0.001). In contrast DAPC results were very similar between datasets, with cluster number, structure and population identity of clusters generally agreeing well (Figure 1, Figure 3c).

*2) M2 Vs. M3*: the effect of reducing the number of samples per population on the inferred population structure. The number of samples per population in the M2 subset (microsatellite data only for the samples used in RADseq, mean = 9.125 ± 0.8) was significantly lower than that of the M3 subset (mean, 19.6 ± 9.0, *t* = -4.66, df = 15, *P* < 0.001), as was the number of alleles per population (M2 mean = 24.4 ± 7.3, M3 mean = 27.4 ± 8.1, *t* = -5.72, df = 15, *P* < 0.001). Population heterozygosities were significantly different between M2 (mean = 0.21) and M3 (mean = 0.23), *t* = -2.4, df = 15, *P* = 0.012), but highly correlated (r = 0.94, *t* = -11.13, *P* < 0.001, df = 15). Pairwise *F*_ST_s were very strongly correlated (r = 0.97, *t* = 46.26, *P* < 0.001, df = 105), but again, still significantly different between the two datasets (M2 mean = 0.46, M3 mean = 0.49, *t* = -6.21, *P* < 0.001, df = 15, Table 4). The patterns of IBD were almost identical for M2 (R^2^ = 0.455, *P* < 0.001) and M3 (R^2^ = 0.447, *P* < 0.001, Supplementary Figure 6) and population structure inferred by DAPC was again similar. BIC scores suggested a similar range of cluster number for M2 and M3, the smallest of which was nine in both cases.

**Table 4.**
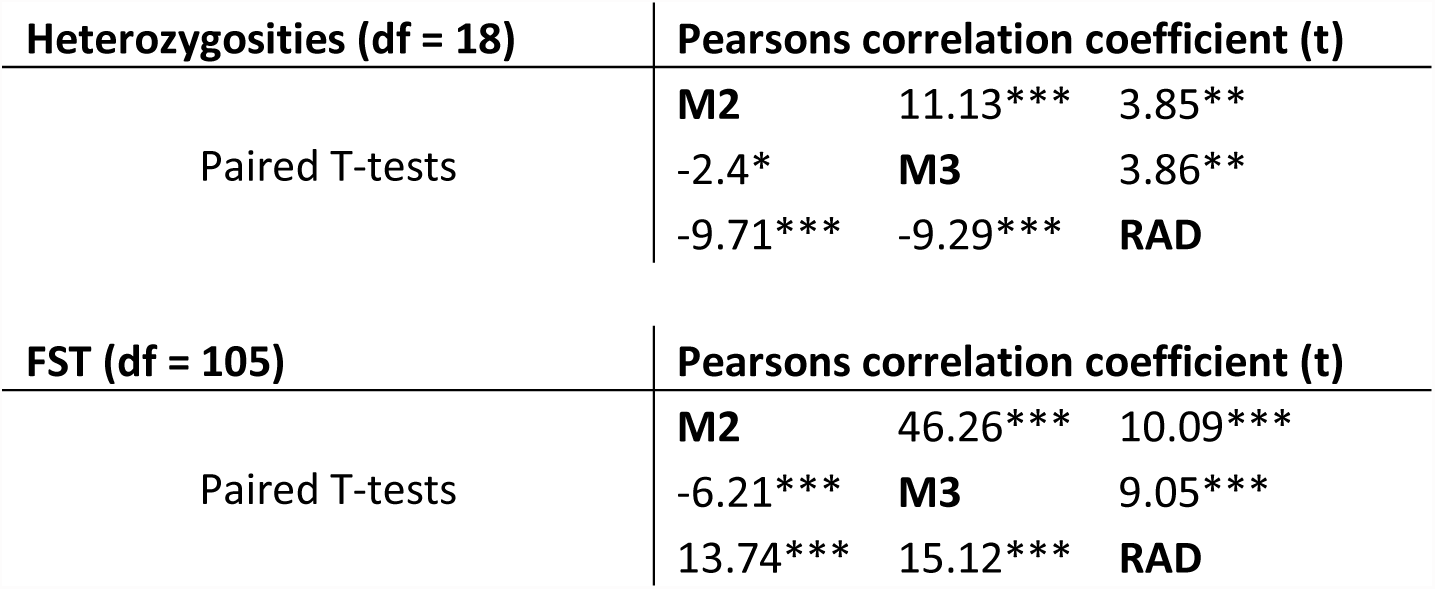
Pearson’s product-moment correlation coefficients and Pared t-tests comparing Heterozygosities and FSTs between M2, M3 and RADseq datasets. *** *P* = <0.001, ** *P* = < 0.005, * *P* = < 0.05.

*3) RADseq Vs. M3:* The effect of the number and the type of markers used on the phylogeographic results. We compared the results from the RADseq and M2 datasets, which contain exactly the same samples (with the exception of three individuals missing in M2). Significant correlations were again found between heterozygosities estimated for the two datasets (r = 0.69, *t* = 3.73, *P* = 0.002, df = 15) and pair-wise *F*_ST_s (r = 0.70, *t* = 10.09, *P* < 0.001, df = 105), but RADseq data yielded much lower pairwise *F*_ST_s (mean RAD = 0.29, mean M2 = 0.46, *t* = 13.74, *P* < 0.001, df = 15). DAPC analysis of RADseq data resolved populations into much more distinct clusters (Figs. 3a, 3b), and the IBD pattern found was considerably stronger in the RADseq (R^2^ = 0.722, *P* < 0.001) dataset compared to M2 (R^2^ = 0.455, *P* < 0.001, Supplementary Figure 6).

## Discussion

In this study, we aimed to simultaneously produce a phylogeographic framework on which to base conservation strategies for *C. carassius* in Europe, and compare the relative suitability of genome-wide SNP markers and microsatellite markers for such an undertaking. Through comparison of the inferred population structure from microsatellite and genome-wide SNP data, we show that there are important differences in the results from each data type, attributable predominantly to marker type, rather than within population sampling or spatial distribution of samples. However, despite these differences, all three data types used (mitochondrial, microsatellite and SNP data) agree that, unlike many other European freshwater fish for which phylogeographic data is available, *C. carassius* has not been able to cross the Danubian catchment boundary into northern Europe. This has resulted in two, previously unknown, major lineages of *C. carassius* in Europe, which we argue should be considered as separate conservation units.

### Phylogeography and postglacial recolonisation of C. carassius in Europe

The most consistent result across all three marker types (mtDNA sequences, microsatellites and RADseq) was the identification of two highly-divergent lineages of *C. carassius* in Europe. The distinct geographic distribution of these lineages; Lineage 1 being widely distributed across north and eastern Europe and Lineage 2 generally only in the River Danube catchment, indicates a long-standing barrier to gene flow between these geographic regions. Bayesian inference based on mtDNA phylogeny and ABC analysis of RADseq data showed remarkable agreement, estimating that these lineages have been isolated for 2.3 MYA (95% CI = 1.30–3.22) and 2.2 (95% CI = 2 – 6.12) MYA respectively, which firmly places the event at the beginning of the Pleistocene (2.6 MYA;(Gibbard & Head 2009). This pattern differs substantially from the general phylogeographic patterns observed in other European freshwater fish. Indeed, previous studies have shown that the Danube catchment has been an important source for the postglacial recolonisation of freshwater fish into northern Europe or during earlier interglacials in the last 0.5 MYA. For example, chub *Leuciscus cephalus (Durand et al. 1999), Eurasian perch Perca fluviatilis (Nesbø et al. 1999), riffle minnow Leuciscus souffia (Salzburger et al. 2003), grayling Thymallus thymallus(Gum et al. 2009), European barbel Barbus barbus (Kotlík & Berrebi 2001), and roach Rutilus rutilus (Larmuseau et al. 2009)* all crossed the Danube catchment boundary into northern drainages such as those of the rivers Rhine, Rhône and Elbe during the mid-to-late Pleistocene. The above species occur in lotic habitats, and most are capable of relatively high dispersal. In contrast *C. carassius* has a very low propensity for dispersal, and a strict preference for the lentic backwaters, isolated ponds and small lakes(Holopainen *et al*. 1997; Culling *et al*. 2006); Copp 1991). We therefore hypothesise that these ecological characteristics of *C. carassius* have reduced its ability to traverse the upper Danubian watershed, which lies in a region characterised by the Carpathian Mountains and the Central European Highlands. This region may have acted as a barrier to the colonisation of *C. carassius* into northern European drainages during the Pleistocene. It should be noted, however, that phylogeography of two species, the spined loach *Cobitis taenia* and European weatherfish *Misgurnus fossilus*, does not support this hypothesis as a general pattern for floodplain species (Janko *et al*. 2005; Culling *et al*. 2006). The former is the only species that we know of other than *C. carassius* showing long-term isolation between the Danube and northern European catchments, but has lotic habitat preferences and good dispersal abilities (Janko *et al*. 2005; Culling *et al*. 2006), whereas the latter inhabits similar ecosystems as *C. carassius*, with low dispersal potential, but has colonised northern Europe from the Danube catchment(Bohlen *et al*. 2006, 2007).

There is one notable exception to the strict separation between Danubian and northern European *C. carassius* populations. The population CZE1, located in the River Lužnice catchment (Czech Republic), which drains into the River Elbe, clusters with Danubian populations in both the microsatellite and mtDNA data. The sample site from the River Lužnice is located in very close proximity to the Danubian catchment boundary and is situated in a relatively low lying area. Therefore some recent natural movements across the watershed between these river catchments, either through river capture events or ephemeral connections, could have been possible. A similar pattern has been shown in some European bullhead *Cottus gobio* populations along the catchment Danube/Rhine catchment border (Riffel & Schreiber 1995). We also observed the presence of two mtDNA haplotypes from Lineage 2 in some individuals from northern German populations (GER1, GER2, GER8), however, one of these haplotypes was shared with Danubian individuals and the results were not confirmed by nuclear markers. Overall this is most likely to be the result of occasional human mediated long-distance dispersal for the purposes of intentional stocking.

Population structure within Lineage 1 is characterised by a pattern of IBD and a loss of allelic richness from eastern to western Europe. This is consistent with the most likely colonisation scenario identified by the DIYABC analysis, indicating a general southeast to northwest expansion from the Ponto-Caspian region towards central and northern Europe (Figure 4). The Ponto-Caspian region, and in particular the Black Sea basin, was an important refugium for freshwater fishes during the Pleistocene glacials, and a similar colonisation route has been inferred for many other freshwater species in northern Europe(Nesbø *et al*. 1999; Durand *et al*. 1999; Culling *et al*. 2006; Costedoat & Gilles 2009). The DIYABC analysis also suggests that there was an interval of > 200 000 years between the split of the Don population (≈ 270 000 years ago) and the next split in the scenario (approx. 33 600 years ago), which marks the main expansion across central and northern Europe. It appears that no further population divergence can be dated back to the time interval between the Riss/Saalian and the Würm/Weichelian glacial periods. This may be because the range of *C. carassius* has not undergone a major change during that time interval, but it is more likely that the signal of expansion during the Riss-Würm interglacial has been eradicated through a subsequent range contraction during the Würm/Weichelian glacial period. The model also suggests that the Würm/Weichelian period was accompanied by a sustained but moderate reduction in population size over almost 9000 years (Bottleneck A, Figure 4), which may reflect general population size reductions during the Riss glaciations or a series of shorter bottlenecks during subsequent range expansion (Ramachandran et al. 2005, Simon et al 2015, Hewitt 2000).

**Figure 4.**
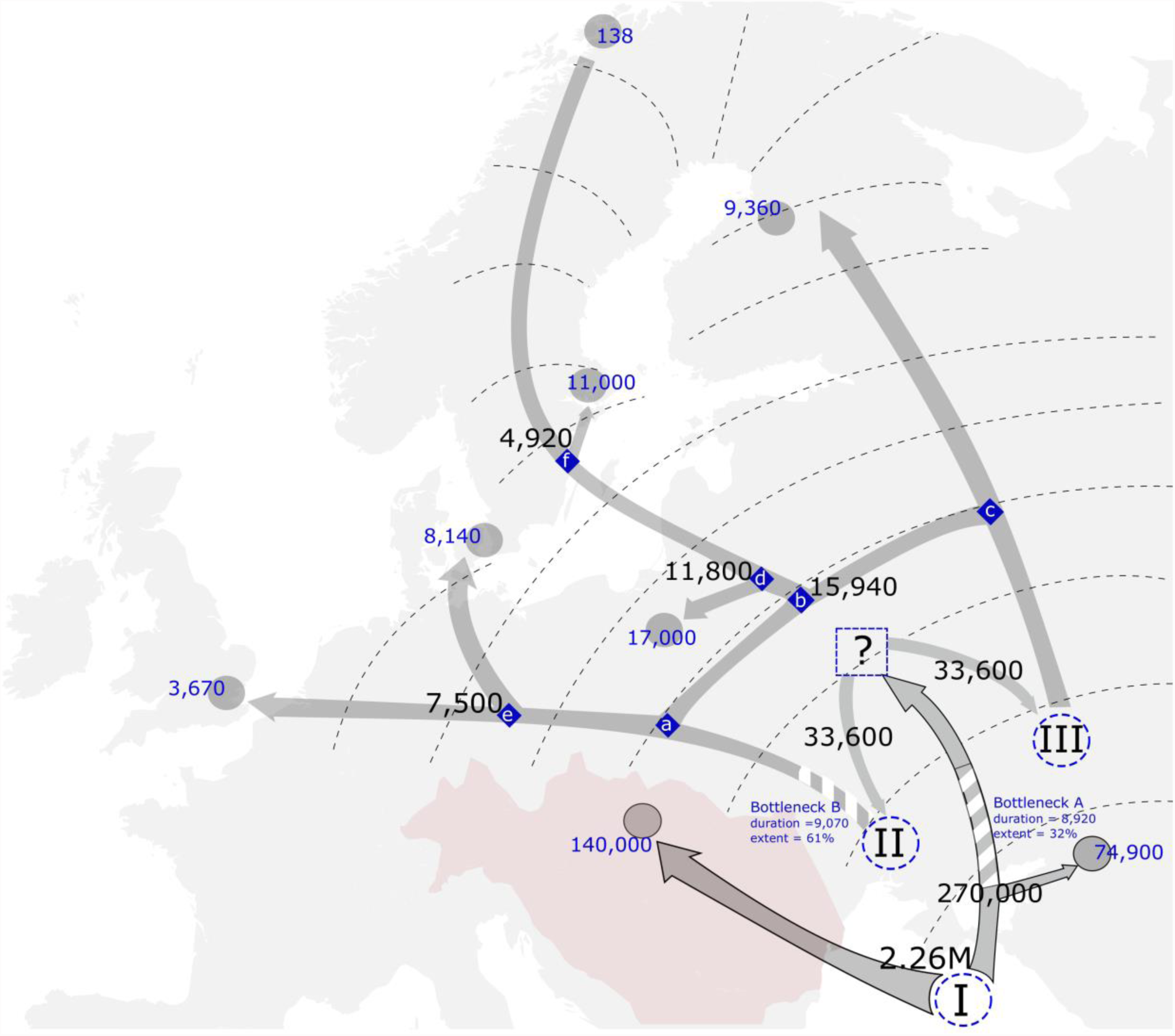
The postglacial recolonisation of C. carassius in Europe. Arrows represent the relationships between population pools used in DIYABC (grey circles) as inferred from Stage 1, scenario 9 (arrows outlined in black) and Stage 3, scenario 14d (arrows with no outline) analyses on RADseq data. Bottlenecks are represented by white-striped sections of arrows. Posterior time estimates in years for each demographic event are given in black, and estimates of Ne are given in blue. Blue diamonds represent ancestral populations inferred by DIYABC and the labels (a-f) correspond to their mention in the text. Hypothetical expansion fronts are represented by dashed contour lines and the Danube river catchment is shaded red. Hypothetical glacial refugia are represented by dashed blue circles (I-III). The blue dashed box (?) represents our inference that C. carassius expanded into central and perhaps northern Europe during the Riss-Würm interglacial, however we cannot estimate this range.

DIYABC analyses inferred the colonisation of northern Europe by two sub-lineages within the mtDNA Lineage 1, which were isolated from each other approximately 33 600 years ago. These sub-lineages may reflect two glacial refugia resulting from the expansion of the Weichselian ice cap to its maximum extent roughly 22 000 years ago (see hypothetical refugia II and III in Figure 4). The western sub-lineage underwent a second long period of population decline (Bottleneck B, Figure 4), which may again represent successive founder effects during range expansion. There is then evidence of secondary contact between these sub-lineages (node b, approximately ≈ 15 940 years ago), contributing to the genetic variation now found in Poland. This inferred admixture event may represent one of the numerous inundation and drainage capture events, which resulted from the melting of the Weichselian ice cap, that are known to have occurred around this time (Grosswald 1980; Gibbard *et al*. 1988; Arkhipov *et al*. 1995). However, as the colonisation of Europe was likely to have occurred via the expansion of colonisation fronts (i.e. dashed contour lines in Figure 4), rather than along linear paths, it could also be indicative of the known IBD gradient between the inferred western and eastern sub-lineages. Such a gradient (eg. between northwestern and northeastern Europe) may give false signals of admixture between intermediate populations, such as those in Poland.

The colonisation of the Baltic sea basin also seems to have been complex, with three independent routes inferred by DIYABC scenario 14d; one recent route through Denmark into southern Sweden, one to the east of the Baltic Sea, through Finland, and one across the Baltic Sea, from populations related to those in Poland (Pool 4). The first of these agrees well with the findings of Janson *et al*.(2014), *whereby populations, including SWE8 from our study (SK3P in Janson et al.(2014)*, in this region were found to be distinct from those in central Sweden. The eastern route shows similarities to the colonisation patterns of *P. fluvilatilis*, which is hypothesised to have had a refugium east of Finland(Nesbø *et al*. 1999) during the most recent glacial period. This is certainly also plausible in *C. carassius* and may account for the distinctiveness of Finnish populations seen in microsatellites and RADseq DAPC analysis. The last colonisation route, across the Baltic Sea from mainland Europe, may have coincided with the freshwater Lake Ancylus stage of the Baltic Sea’s evolution, which existed from ≈ 10 600 to 7 500 years ago (Björck 1995; Kostecki 2014). The Lake Ancylus stage likely provided a window for the colonisation of many of the species now resident in the Baltic, and has been proposed as a possible window for the colonisation of *Thymallus thymallus(Koskinen et al. 2000), Cobitis taenia*,(Culling *et al*. 2006), *Cottus gobio(Kontula & Väinölä 2001)* and four *Coregonus* species(Svärdson 1998). Consistent with this, we found strong similarity between populations from Fasta Åland, southern Finland and central Sweden, suggesting that shallow regions in the central part of Lake Ancylus (what is now the Åland Archipelago), may have provided one route across Lake Ancylus.

It is also likely that the contemporary distribution of *C. carassius* in the Baltic has been influenced by human translocations. *C. carassius* were often used as a food source in monasteries in many parts of Sweden(Janson *et al*. 2014), and the Baltic island of Gotland(Rasmussen 1959; Svanberg *et* al. 2013) was an important trading port of the Hanseatic League – a commercial confederation that dominated trade in northern Europe from the 13^th^ to 17^th^ centuries. Previous data suggest that *C. carassius* was transported from the Scania Province, southern Sweden, where *C. carassius* aquaculture was common at least during the 17^th^ century, to parts further north (Svanberg *et al*. 2013; Janson *et al*. 2014).

### Implications for the conservation of *C. carassius* in Europe

The two *C. carassius* lineages exhibit highly-restricted gene flow between them and are the highest known organisational level within the species. They therefore meet the genetic criteria for Evolutionarily Significant Units (ESUs) as described in (Fraser & Bernatchez 2001). This is especially important in light of the current *C. carassius* decline in the Danubian catchment (Bănărescu 1990; Navodaru *et al*. 2002; Lusk *et al*. 2010; Savini *et al*. 2010). The conservation of *C. carassius* in central Europe must therefore take these catchment boundaries into consideration, as opposed to political boundaries. A first step would be to include *C. carassius* in Red Lists, not only for individual countries, but at the regional (e.g. European Red List of Freshwater Fishes; (Freyhof & Brooks 2011) and global (IUCN 2014) scales, and we hope that the evidence presented here will facilitate this process. Within the northern European lineage, the Baltic Sea basin shows high levels of population diversity, likely owing to its complex colonisation history. As such, the Baltic represents an important part of the *C. carassius* native range. Although *C. carassius* is not currently thought to be threatened in the Baltic region, *C. gibelio* is invading this region and is considered a threat (Urho & Lehtonen; Deinhardt 2013).

### Microsatellites vs RADseq for phylogeography

Broad conclusions drawn from each of our RADseq-derived SNPs, full or partial microsatellite datasets are consistent, demonstrating deep divergence between northern and southern European populations and an IBD pattern of population structure in northern Europe. However, two striking differences exist in the phylogeographic results produced by RADseq compared to those of the microsatellite datasets. Firstly, the IBD pattern inferred from RADseq data was considerably stronger than for any of the microsatellite datasets. This effect was also found by Coates *et al*. (2009) when comparing SNPs and microsatellites, who postulated that it was driven by the differences in mutational processes of the markers. The second major difference between RADseq and microsatellite results was that clusters inferred by DAPC from the RADseq data were considerably more distinct compared to the full microsatellite dataset, emphasising the fine scale structure in the data (which is particularly apparent in the northern Finnish populations). We ruled out the possibility of these differences being caused by the reduction in number of populations, their spatial uniformity or number of individuals per population used in RADseq by creating two partial microsatellite datasets and comparing these to results from the RADseq-SNPs. Differences between marker types were consistently reproducible whether full or partial microsatellite datasets were used in the analyses.

It is also worth noting that the number of populations or the number of samples per population had no apparent impact on IBD and DAPC results between the microsatellite datasets. This is in contrast to predictions of patchy sampling of IBD made by Schwartz and McKelvey (2009), perhaps because of the strong population structure in *C. carassius,* and likelihood that a sufficiently informative number of populations was included even in the reduced datasets.

### Conclusions

We have identified the most likely routes of post-glacial colonisation in *C. carassius*, which deviate from the general patterns observed in other European freshwater fishes. This has resulted in two, previously-unidentified major lineages in Europe, which future broad-scale monitoring and conservation strategies should take into account.

Although our RADseq sampling design included only 17.6% of samples included in the full microsatellite dataset this was sufficient to produce a robust phylogeography in agreement with the microsatellite dataset, and emphasised the fine scale structure among populations. We therefore conclude that RADseq would present the better option for the phylogeography of *C. carassius,* with the huge number of SNP loci overcoming the limitations imposed by reduced sample number.

## Acknowledgments

The authors thank the FSBI (www.fsbi.org.uk) and Cefas (Lowestoft, UK) for funding this research. We thank the following landowners, and contributors of fish tissue; Keith Wesley, Ian Patmore and Dave Emson (England), L. Urho (Helsinki, Finland), M. Himberg (Salo, Finland), J. Krekula (Steninge Castle, Uppland, Sweden), B.-M. Josephson and G. Josephson (Styrstad Vicarage, Sweden), G. Hellström (UmeÅ University, Sweden), K.Ø. Gjelland (NINA, Tromsø, Norway), N. Hellenberg (Gotland Island, Sweden), A. Tuvikene (Center for Limnology, Tartu, Estonia), S.V. Mezhzherin (Kiev, Ukraine), K. Lindström (Kvicksund, Sweden), K.-J. Dahlbom and G. Sundberg (Åland Island, Finland), O. Sandström and M. Andersson (Skutab, Öregrund, Sweden), B. Tengelin (Structor Miljöteknik AB), A. Olsén-Wannefjord (Uppsala, Sweden), Müller Tamás (Godollo, Hungary), András Weiperth (Hungary), Peter D Rask Møller and Henrik Carl (Copenhagen, Denmark), Oksana Stoliar, (Ternopil, Ukraine), Manuel Deinhardt (Jyväskylä, Finland).

## Supplementary materials

### Detecting hybrids

#### Methods

In total we acquired tissue samples of 1078 Fish during sampling for this study. All of which were first genotyped using multiplex 1 (Supplementary table 1) which contained the 6 species diagnostic microsatellite loci. These data were then analysed using the NewHybrids v. 1.1 (Anderson & Thompson 2002) software package in order to determine whether each fish was *C. carassius, C. a. auratus, C. a. Gibelio* or a hybrid between any of these species.

NewHybrids uses allele frequencies to give a likelihood probability that an individual belongs to one species or another, or if the individual one of several hybrid classes (F1, F2 or backcross). Data from 20 *C. carassius* samples, which were confidently identified as pure from both morphology and genotypes, and were not sympatric with non-native species, were included in each analysis as baseline data. Priors were then added to the analyses specifying that these individuals were indeed pure in order to give the software more power with which to assess allele frequencies associated with *C. carassius*. To be sure to account for allele frequency differences between different geographic regions, only pure individuals from regions neighbouring the hybrid population were used. Individuals which had more than a 25% chance of being an F1 hybrid, F2 hybrid, or a backcross were removed from population structure analyses and were not genotyped at the additional 7 microsatellite loci (Multiplexes 2.1 and 2.2, Supplementary table 1).

#### Results

Of the 1087 genotyped fish from 58 populations, 942 individuals across 55 populations (86.7%) were identified as pure crucian using the first set of 6 species diagnostic loci in NewHybrids analyses. 19(1.8%) from 2 different populations were identified as *C. a. auratus*, 15 fish (1.4%) from 4 populations were identified as. *C. a. gibelio* and 10 fish (0.93%) from two populations were identified as *C. carpio*. NewHybrids identified 60(5.5%) *C. carassius × C. a. auratus* hybrids, 25(2.2%) *C. carassius × C. a. gibelio* hybrids, and 16(1.5%) *C. carassius × C. carpio* hybrids. Of the 942 fish identified as pure *C. carassius*, 848 in 49 populations existed in sites where hybrids or non-native species were not detected by microsatellite genotyping. To safeguard against cryptic introgression which may produce erroneous results only these 848 pure *C. carassius* were used for the main phylogeographic analyses and tests of the status of *C. carassius* in England.

### DAPC & Running parameters

#### Methods

Population structure was examined using Discriminant Analyses of Principal Components (DAPC, (Jombart *et al*. 2010)) in adegenet. Similar to the more commonly used program, STRUCTURE (Pritchard et al. 2000), DAPC is an individual-based approach that uses Principal Components Analysis (PCA) to transform population genetic data and Discriminant Analysis (DA) to identify clusters. The number of clusters is assessed using the K-means method, which is also used in STRUCTURE (Pritchard *et al*. 2000). Unlike STRUCTURE, DAPC does not assume underlying population genetics models such as Hardy-Weinberg Equilibrium (Jombart *et al*. 2010) and is therefore more suitable for analysing C. carassius since populations are often bottlenecked (Hänfling *et al*. 2005). An additional benefit of DAPC is that it maximizes between-group variation, while minimizing variation within groups, allowing for optimal discrimination of between-population structure (Jombart *et al*. 2010).

#### Results

For the full microsatellite dataset (M1), BIC scores indicated that between 11 and 19 genetic clusters (Supplementary Figure 5) would be an appropriate model of the variation in the data. We therefore chose 11 clusters to use in the discriminant analysis, retaining 8 principal components as recommended by the spline interpolation a-scores (Supplementary Figure 5c) and we kept 2 linear discriminants for plotting (Figure 1b).

Three major lineages were found, one located in the Danube, one in the Don, and one spread across northern Europe. However the large amount of diveregence between them masked the population structure present in northern Europe. We therefore subsetted the data, separating NEU populations from RUS1, GER3, GER4, CZE1 (and SWE9, which was an outlier within NEU, Figure 1b) and reanalysed them with DAPC in order to better infer fine population structure between them.

For the RADseq dataset, BIC scores suggested between 9 and 14 genetic clusters, similar to the range inferred in the microsatellite data, we therefore chose 9 clusters to take forward in the analysis. As recommended by spline interpolation, we retained 7 principal components and we kept 2 of the linear discriminants from the subsequent discriminant analysis

### Assessment of spatial uniformity of sampling locations

#### Methods

In order to assess the geographic uniformity of the sampling regimes in each data subset, we used two measures of spatial patterns. The nearest neighbour distance distribution function (G), measures the distance of each sampling location to its nearest neighbour (Ripley 1991). The L-function is a transformation (for ease of interpretation) of Ripley’s K-function (Ripley 1991), which measures the number of sampling locations within a given radius from each point. K has the advantage of assessing the uniformity of the sampling regime over multiple scales, as opposed to only measuring distances between closest neighbours as with G. In both cases, the estimates of G or K from our sampling locations were compared against random poisson distributions, which would represent uniformly spaced sampling locations. 5% and 95% confidence thresholds for these poisson distributions were also calculated to allow us to determine whether our sampling regimes significantly deviated from random (p <0.05). These calculations were performed using the Gest and Lest functions (for G and L respectively) in the package “spatstats” in R (Baddeley & Turner 2005)

#### Results

Both methods used for the assessment of geographic uniformity of sampling locations shows that the M1 dataset locations are more patchily distributed than those of the M2, M3 and RAD datasets (Supplementary Figure 10).

### Additional discussion

#### Population structure in northwest Europe

An intriguing result lies in the genetic similarity between populations in England with those in Belgium and Germany. *C. carassius* has been designated as native to England, however this status has been contentious in the past (Maitland 1972). Under the assumption that it is native, and considering the observed diversity and divergence times between populations across mainland Europe, we would expect to see stronger population structure between English and continental Europe, which have been separated for approximately 7800 years (Coles 2000). Given the observed diversity between populations across mainland Europe, which, according to DIYABC analysis, has arisen relatively recently. Clearly further examination of this issue is warranted and molecular data would be a value addition to the current evidence, which is predominantly anecdotal.

**Supplementary table 1.** Microsatellite loci used, grouped by their combinations in multiplex reactions. Multiplex primer mix ratios for PCR were chosen so as to give even peak strengths when analysing PCR products. Allele size ranges are those present in C. carassius for all 43 putatively pure crucian populations.

| Locus | Multiplex # | Primer mix Ratios* | # Alleles | Allele size range | Ho | GenBank Accession no. | Reference |
| --- | --- | --- | --- | --- | --- | --- | --- |
| GF1 | 1 | 0.1 | 1 | 299 | 0 | U35614 | Zheng et al. 1995 |
| GF17 | 1 | 0.1 | 2 | 182-186 | 0.024 | U35616 | Zheng et al. 1995 |
| GF29 | 1 | 0.2 | 8 | 191-226 | 0.348 | U35618 | Zheng et al. 1995 |
| J7 | 1 | 0.07 | 10 | 202-228 | 0.109 | AY115095 | Yue & Orban 2002 |
| MFW2 | 1 | 0.1 | 1 | 161 | 0 | - | Croojimans et al. 1997 |
| Ca07 | 1 | 0.2 | 9 | 122-140 | 0.286 | D85428 | Yue & Orban 2004 |
| TE Buffer | 1 | 0.23 |  |  |  |  |  |
| J69 | 2.1 | 0.4 | 14 | 213-241 | 0.404 | AY115106 | Yue & Orban 2002 |
| HJLY17 | 2.1 | 0.1 | 9 | 152-168 | 0.223 | DQ378986 | Zhi-Ying et al. 2006 |
| HJLY35 | 2.1 | 0.1 | 18 | 261-307 | 0.377 | DQ403242 | Zhi-Ying et al. 2006 |
| TE Buffer | 2.1 | 0.4 |  |  |  |  |  |
| J20 | 2.2 | 0.2 | 9 | 171-218 | 0.149 | AY115099 | Yue & Orban 2002 |
| J58 | 2.2 | 0.1 | 14 | 119-147 | 0.398 | - | Yue & Orban 2002 |
| MFW7 | 2.2 | 0.35 | 25 | 160-206 | 0.464 | - | Croojimans et al. 1997 |
| MFW17 | 2.2 | 0.35 | 26 | 185-262 | 0.41 | - | Croojimans et al. 1997 |

\* All primers used at 10mM per ul concentration, diluted in ddH2O from 100mM per ul stock

**Supplementary table 2.** Haplotype memberships for 101 Cytochrome B sequences used in Fig. 2.

|  | Haplotype | N | Drainage<br>(n populations) | Sample<br>sequence |
| --- | --- | --- | --- | --- |
| Lineage 1 | Hap 1 | 3 | Baltic | FIN5 1-3 |
|  | Hap 2 | 1 | Baltic | EST1 2 |
|  | Hap 3 | 49 | Elbe(2), Baltic(24),<br>Scheldt(1),<br>Rhine(2), North<br>sea(2), Vistula(6),<br>Volga(4), Don(3),<br>Danube(1),<br>Hunte(4) | GER1 1,3, EST1 1, 3, SWE6 1 -3, BEL1 3 , GER2 2, 3, GER4 2, NOR 1, 2,<br>SWE11 1-3, RUS2 2 , RUS4 1, 3 , FIN1 1-3, FIN4 1-3, POL4 1-3, RUS1 1-3,<br>SWE8 1-3, POL5 1-3, SWE4 1-3, RUS3 1, 3, 4, CZE2 1, GER6 1 – 4, SWE14 1, SWE15<br>1 |
|  | Hap 4 | 1 | Volga | RUS2 1 |
|  | Hap 5 | 1 | Baltic | RUS4 2 |
|  | Hap 6 | 1 | Dnieper | BLS 3 |
|  | Hap 7 | 1 | Volga | RUS3 2 |
|  | Hap 8 | 3 | Baltic | SWE3 1-3 |
|  | Hap 9 | 2 | Baltic | SWE2 1, 2 |
|  | Hap 10 | 1 | Baltic | SWE2 3 |
|  | Hap 11 | 3 | Baltic | SWE9 1-3 |
|  | Hap 12 | 13 | UK(4), Rhine(1),<br>Baltic (2) | GBR7 1, GBR6 1-3, GBR8 1-3, NET 1, GER5 1-3, GBR12 1, 2 |
|  | Hap 13 | 3 | Baltic | FIN3 1-3 |
| Lineage 2 | Hap 14 | 3 | Danube | GER4 1, 2, AUS3<br>1 |
|  | Hap 15 | 3 | Elbe(1), Rhine(1),<br>Danube(1) | GER1 2, GER2 1, AUS2 1 |
|  | Hap 16 | 1 | Danube | CZE1 1 |
|  | Hap 17 | 1 | Danube | CZE1 2 |
|  | Hap 18 | 1 | Danube | CZE1 3 |
|  | Hap 19 | 2 | Danube | HUN 1, 2 |
|  | Hap 20 | 3 | Danube | GER3 1-3 |
|  | Hap 21 | 2 | Danube | AUS1 1, 2 |
|  | Hap 22 | 2 | Lahn | GER7 1, 2 |

**Supplementary table 3.**
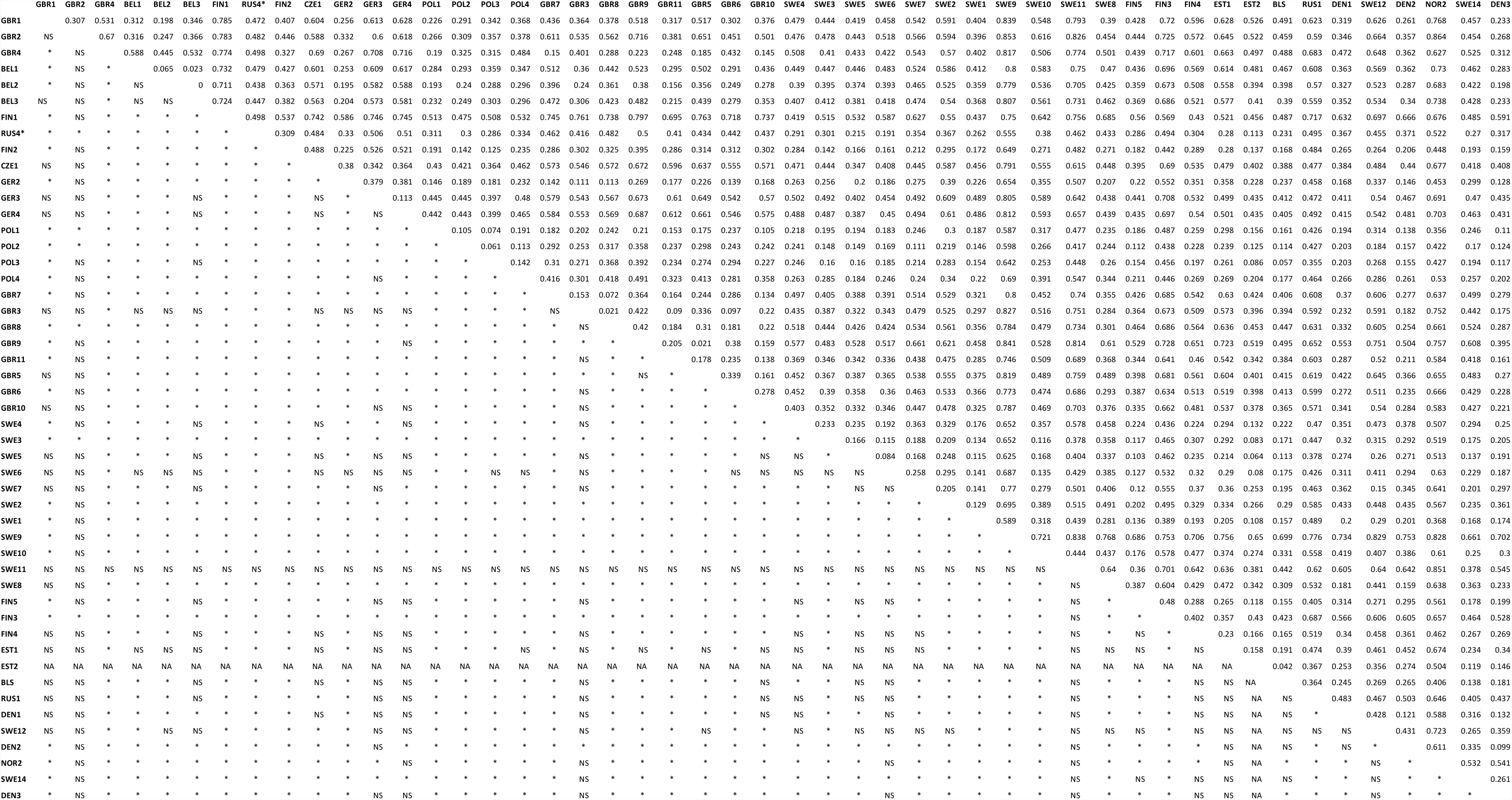
Pairwise FST values calculated using the M1 dataset.

**Supplementary table 4.**
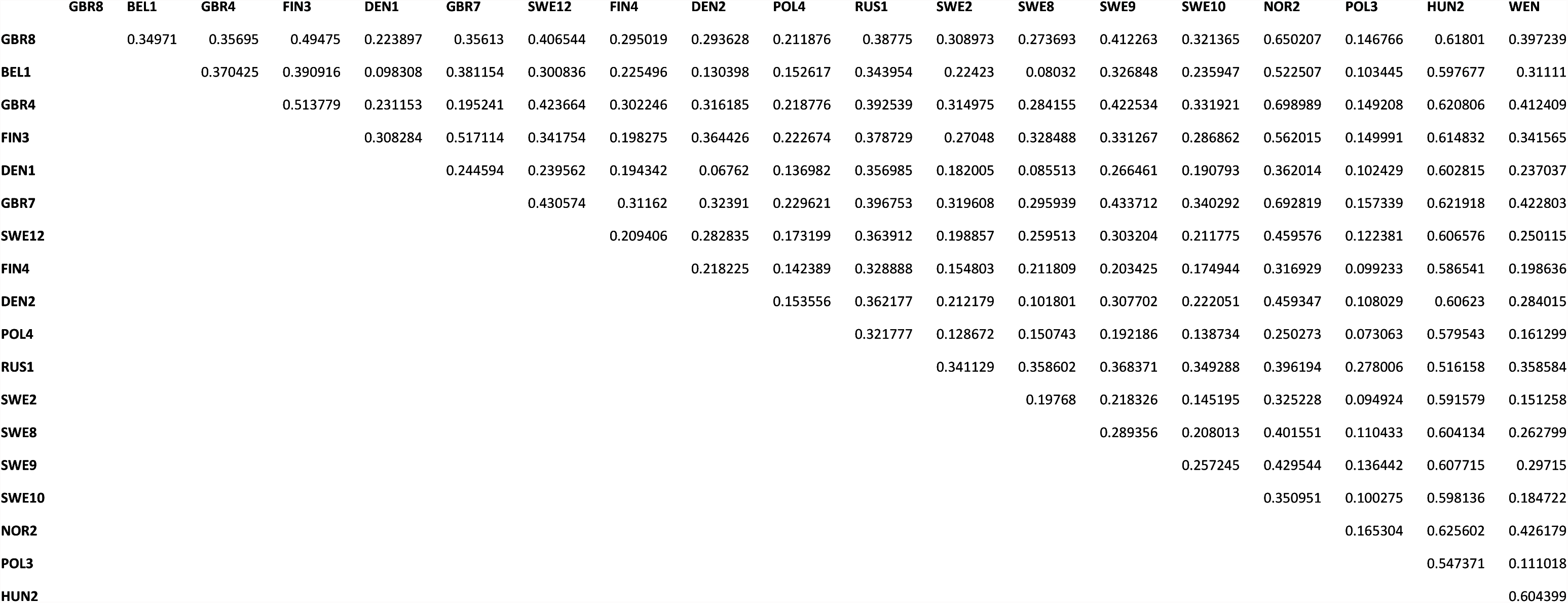
Pairwise FST values calculated using the RADseq dataset.

**Supplementary table 5.** Genbank accession numbers for the mtDNA sequences used in this study.

| Sample code | Accession number |
| --- | --- |
| FIN5_01 | KT630314 |
| FIN5_02 | KT630315 |
| FIN5_03 | KT630316 |
| EST1_02 | KT630317 |
| GER1_01 | KT630318 |
| EST1_01 | KT630319 |
| GER1_03 | KT630320 |
| SWE6_01 | KT630321 |
| SWE6_02 | KT630322 |
| SWE6_03 | KT630323 |
| BEL1_03 | KT630324 |
| EST1_03 | KT630325 |
| GER2_02 | KT630326 |
| GER2_03 | KT630327 |
| GER4_02 | KT630328 |
| NOR1_01 | KT630329 |
| NOR1_02 | KT630330 |
| SWE11_01 | KT630331 |
| SWE11_02 | KT630332 |
| SWE11_03 | KT630333 |
| RUS2_02 | KT630334 |
| RUS4_01 | KT630335 |
| RUS4_03 | KT630336 |
| FIN1_01 | KT630337 |
| FIN1_02 | KT630338 |
| FIN1_03 | KT630339 |
| FIN4_01 | KT630340 |
| FIN4_02 | KT630341 |
| FIN4_03 | KT630342 |
| POL4_01 | KT630343 |
| POL4_02 | KT630344 |
| POL4_03 | KT630345 |
| RUS1_01 | KT630346 |
| RUS1_02 | KT630347 |
| RUS1_03 | KT630348 |
| SWE8_01 | KT630349 |
| SWE8_02 | KT630350 |
| SWE8_03 | KT630351 |
| POL3_01 | KT630352 |
| POL3_02 | KT630353 |
| POL3_03 | KT630354 |
| SWE4_01 | KT630355 |
| SWE4_02 | KT630356 |
| SWE4_03 | KT630357 |
| RUS3_01 | KT630358 |
| RUS3_03 | KT630359 |
| RUS3_04 | KT630360 |
| RUS2_01 | KT630361 |
| RUS4_02 | KT630362 |
| BLS_03 | KT630363 |
| RUS3_02 | KT630364 |
| SWE3_01 | KT630365 |
| SWE3_02 | KT630366 |
| SWE3_03 | KT630367 |
| SWE2_01 | KT630368 |
| SWE2_02 | KT630369 |
| SWE2_03 | KT630370 |
| SWE9_01 | KT630371 |
| SWE9_02 | KT630372 |
| SWE9_03 | KT630373 |
| GBR7_01 | KT630374 |
| GBR6_01 | KT630375 |
| GBR8_01 | KT630376 |
| GBR8_02 | KT630377 |
| GBR8_03 | KT630378 |
| GBR6_02 | KT630379 |
| GBR6_03 | KT630380 |
| CZE1_01 | KT630381 |
| CZE1_02 | KT630382 |
| CZE1_03 | KT630383 |
| GER4_01 | KT630384 |
| GER4_03 | KT630385 |
| GER1_02 | KT630386 |
| GER2_01 | KT630387 |
| FIN3_01 | KT630388 |
| FIN3_02 | KT630389 |
| FIN3_03 | KT630390 |
| HUN1_02 | KT630391 |
| GER3_01 | KT630392 |
| GER3_02 | KT630393 |
| GER3_03 | KT630394 |

**Supplementary Figure 1.**
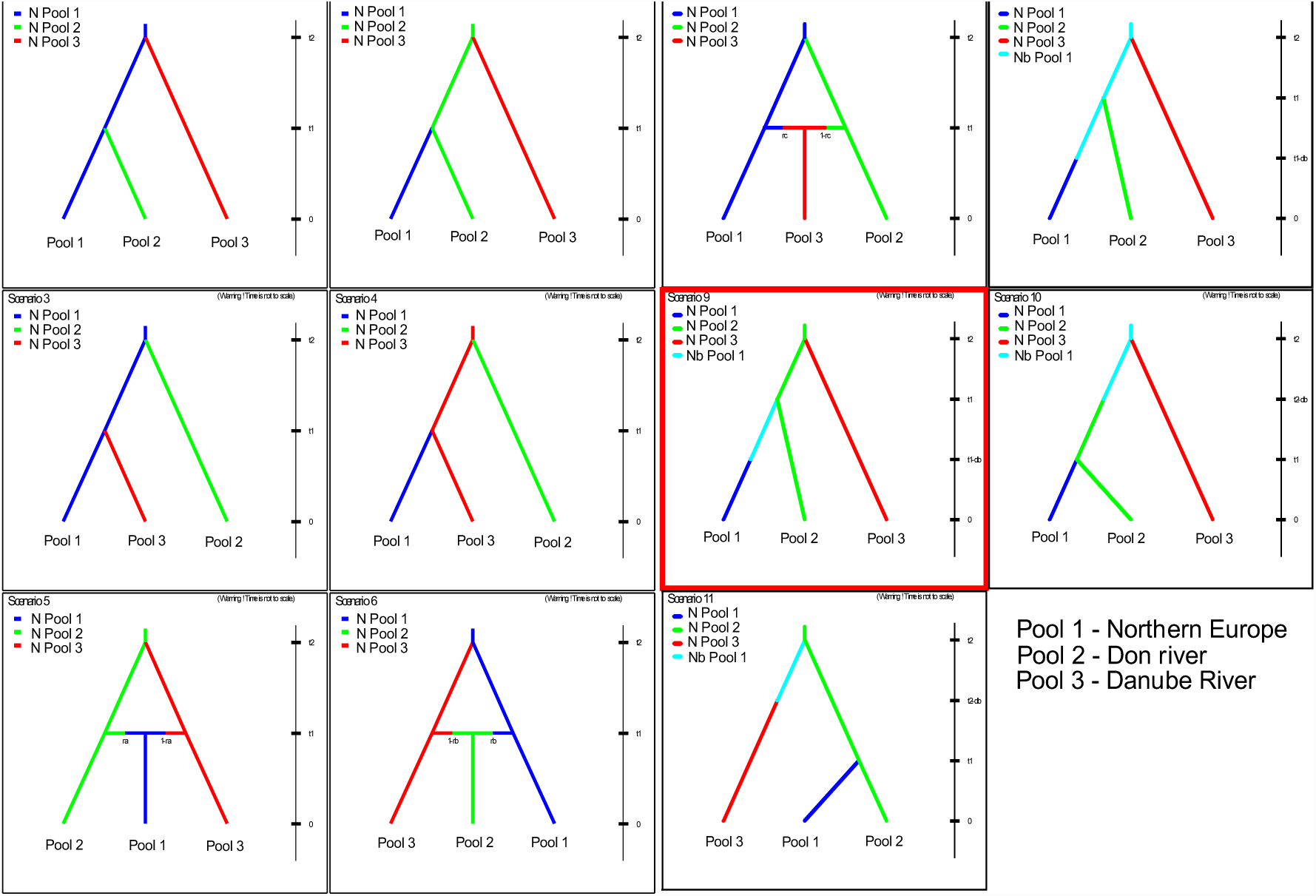
DIYABC scenarios used in broad-scale analysis (Stage 1). See text for population poolings. See Table 3 for population poolings and prior parameter values.

**Supplementary Figure 2.**
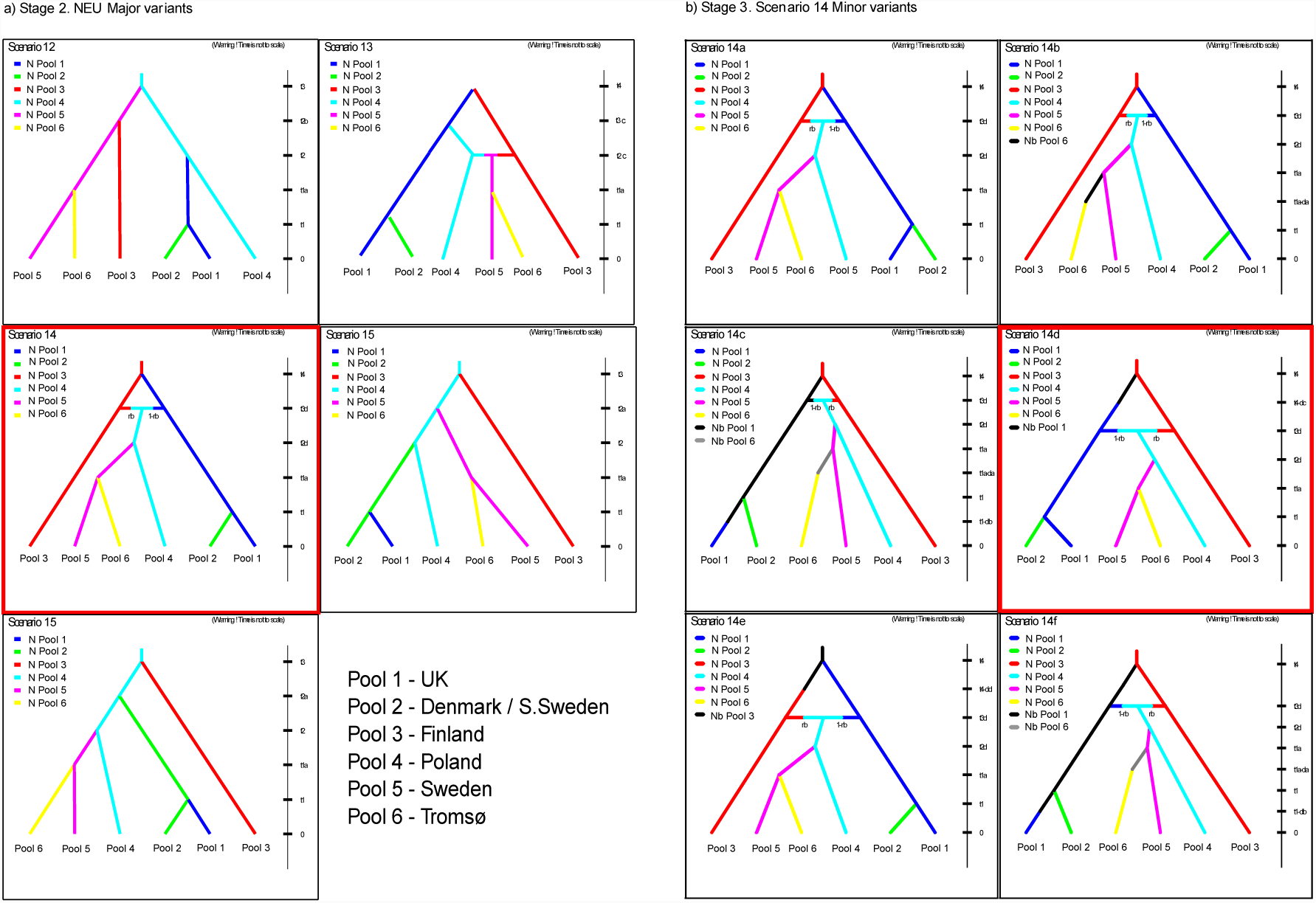
All scenarios tested in stage 2 **a)** and stage 3 **b)** of DIYABC analysis. See Table 3 for population poolings and prior parameter values.

**Supplementary Figure 3.**
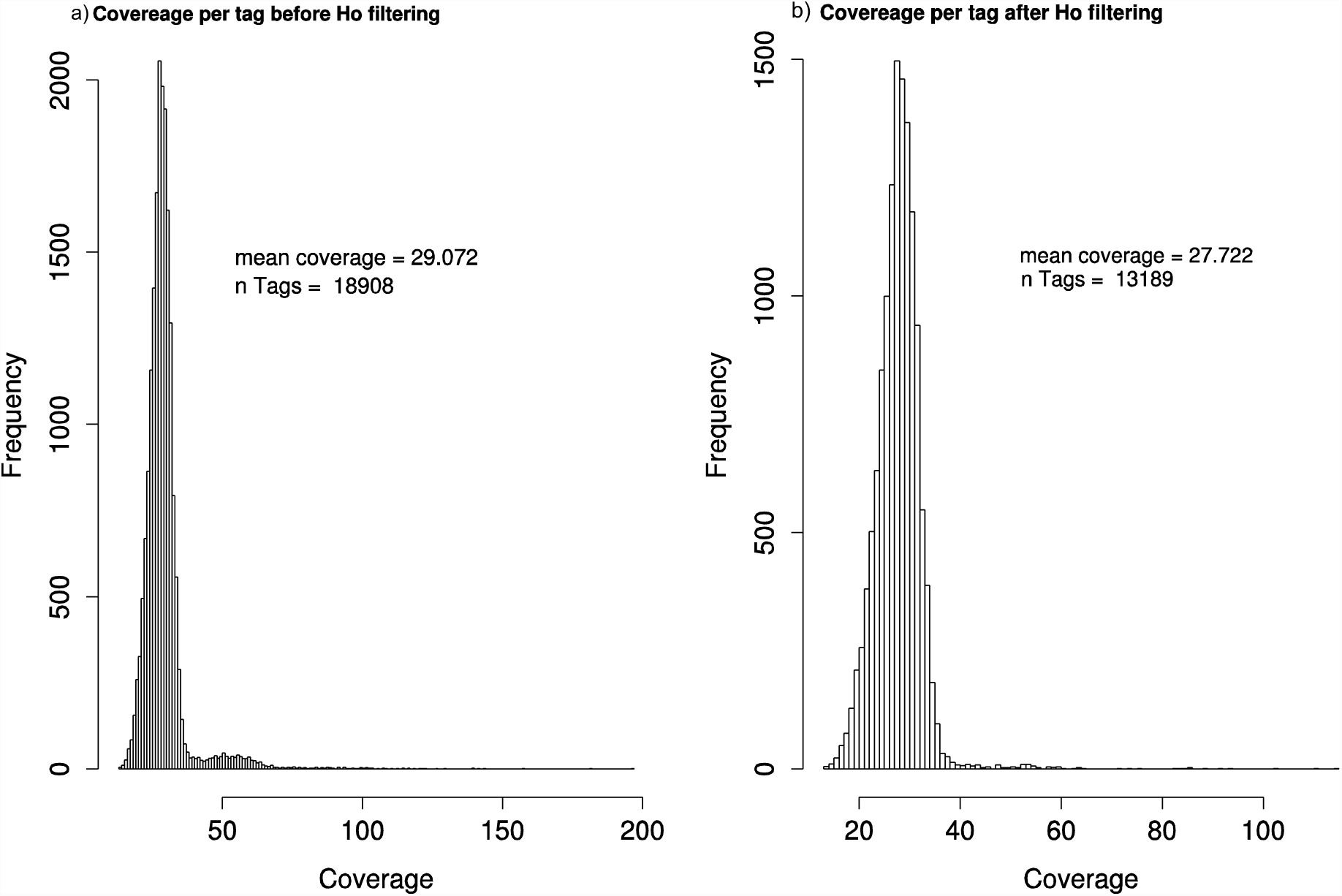
Filtering out merged ohnologs. **a)** Distribution of SNP locus coverage prior to removing loci that had observed heterozygosity higher than 0.5 in one or more population. **b)** Distribution of locus coverage after filtering, showing a loss of many high coverage loci and a reduction in mean SNP coverage. Note the loss of loci with high coverage.

**Supplementary Figure 4.**
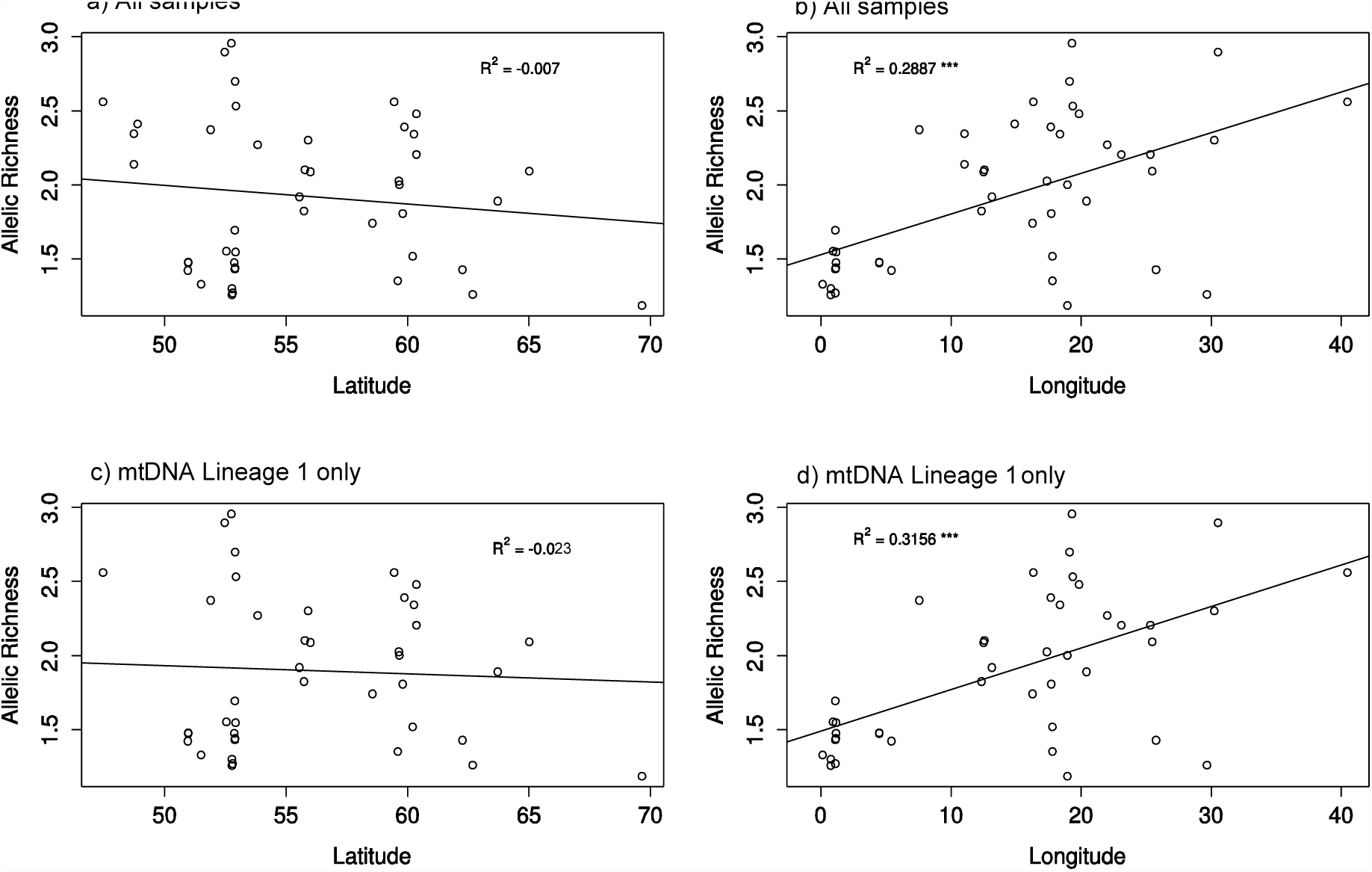
Linear regressions for all samples **a)** Ar against latitude; **b)** Ar against longitude and for only samples in mtDNA lineage 1 **c)** Ar against latitude; d) Ar against longitude.

**Supplementary Figure 5.**
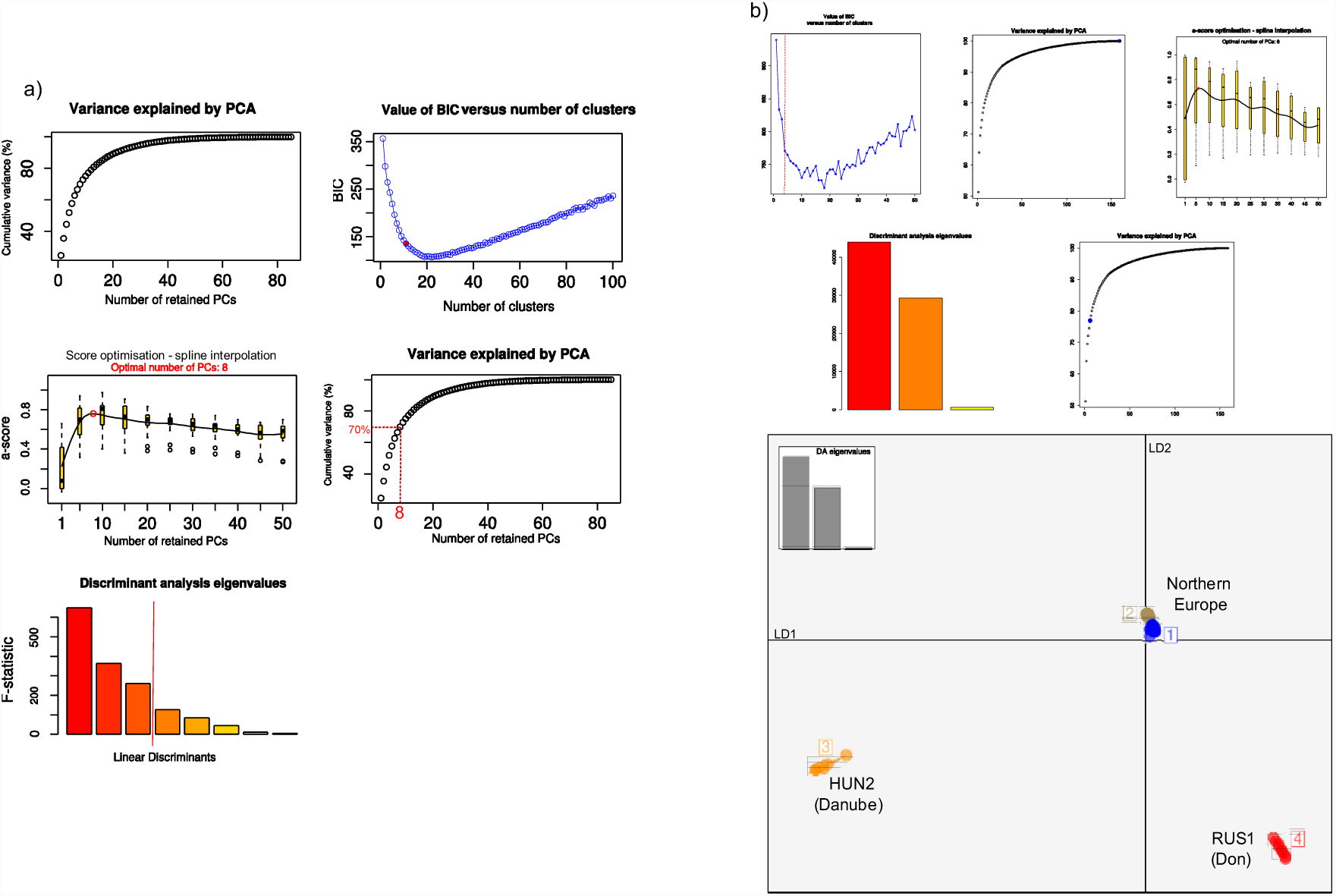
DAPC analysis of **a)** full microsatellite dataset (Excluding NOR2); for results used in Fig. 1) and **b)** Full RADseq dataset.

**Supplementary Figure 6.**
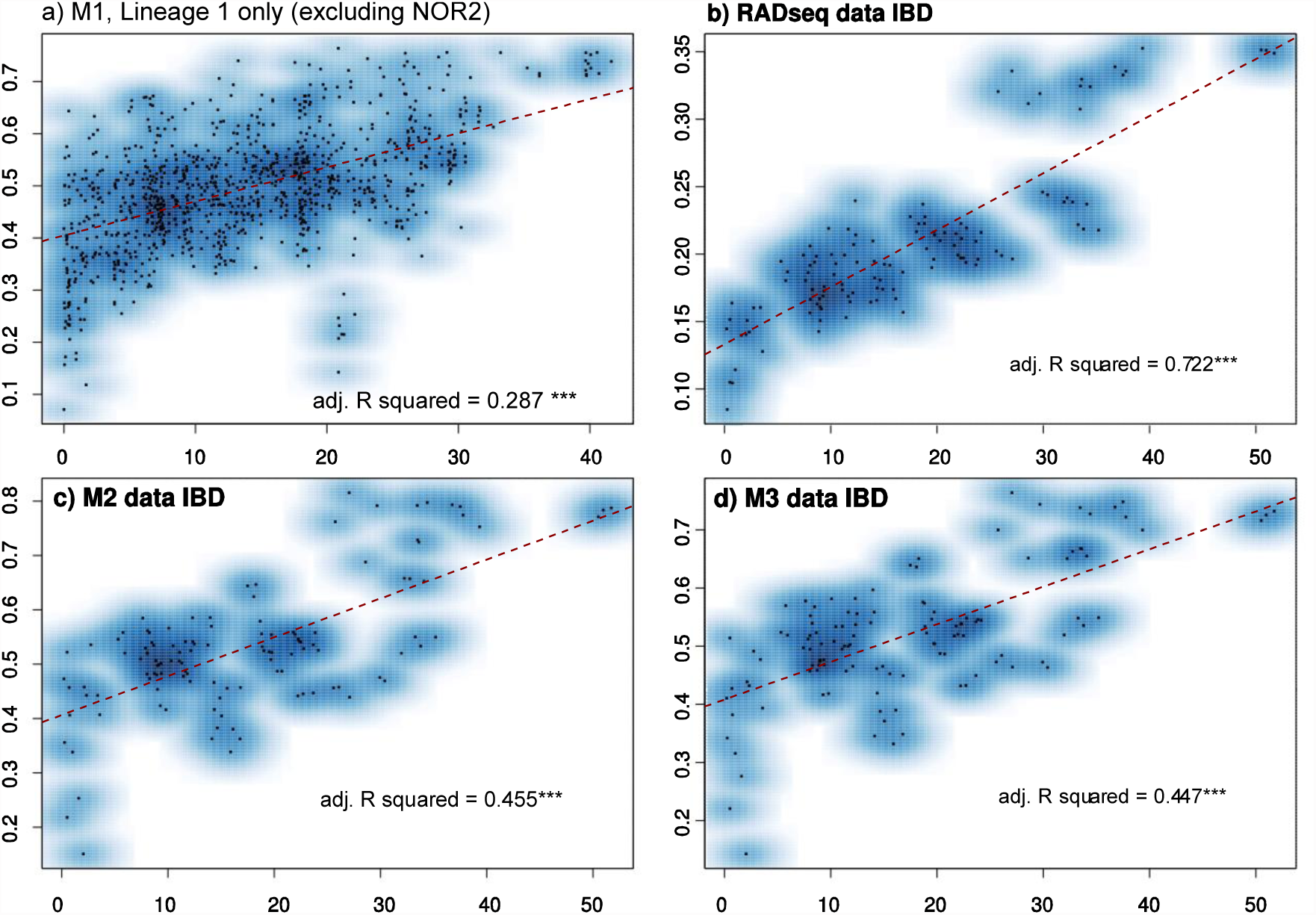
Isolation by distance **a)** in M1 dataset for mtDNA lineage 1 only (excluding NOR2), **b)** Full RADseq dataset, **c)** M2 dataset and **d)** M3 dataset.

**Supplementary Figure 7.**
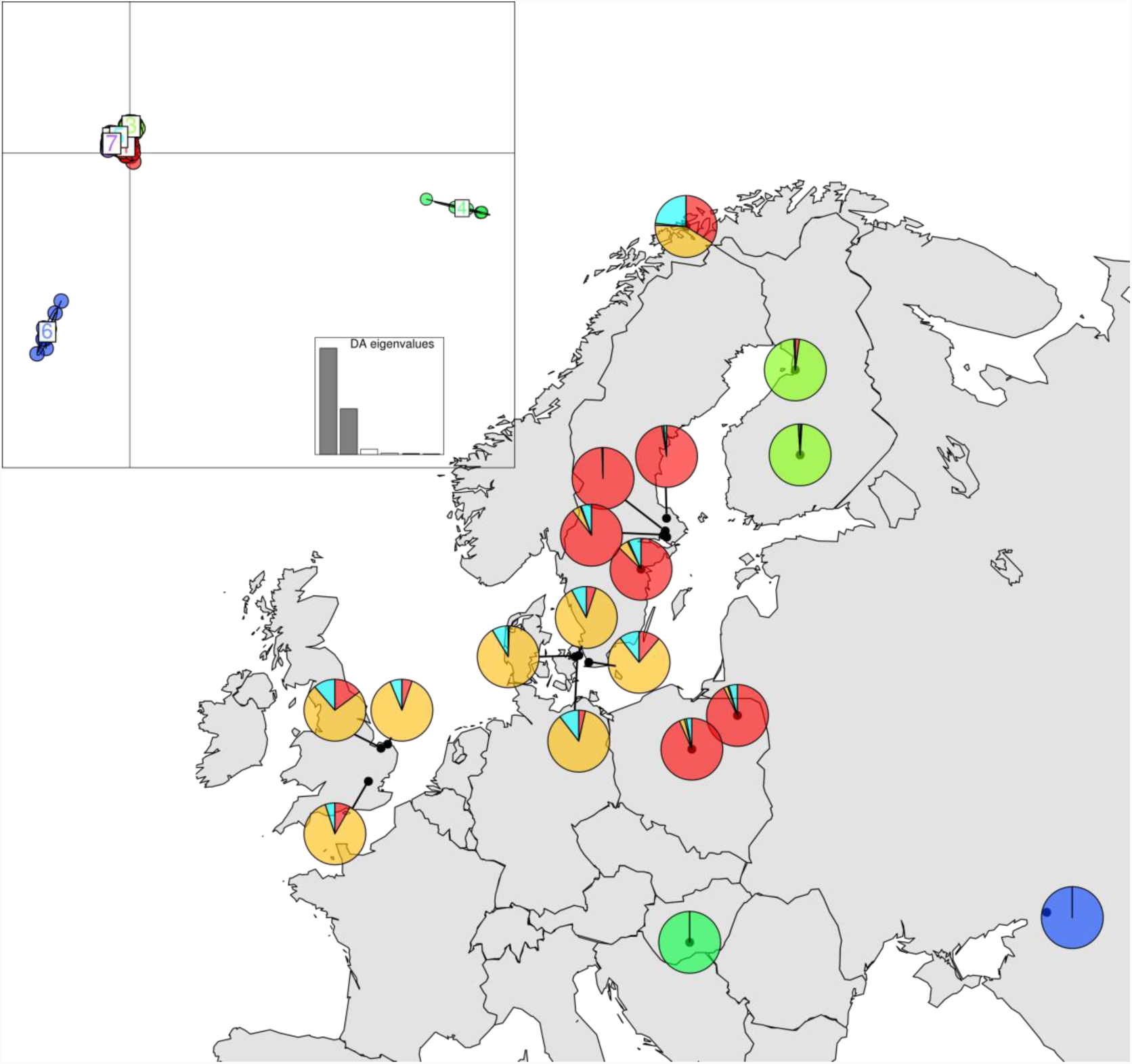
DAPC scatter plot for the 100 SNP RADseq dataset used in the DIYABC analysis, showing the same population structure as inferred from the full RADseq dataset.

**Supplementary Figure 8.**
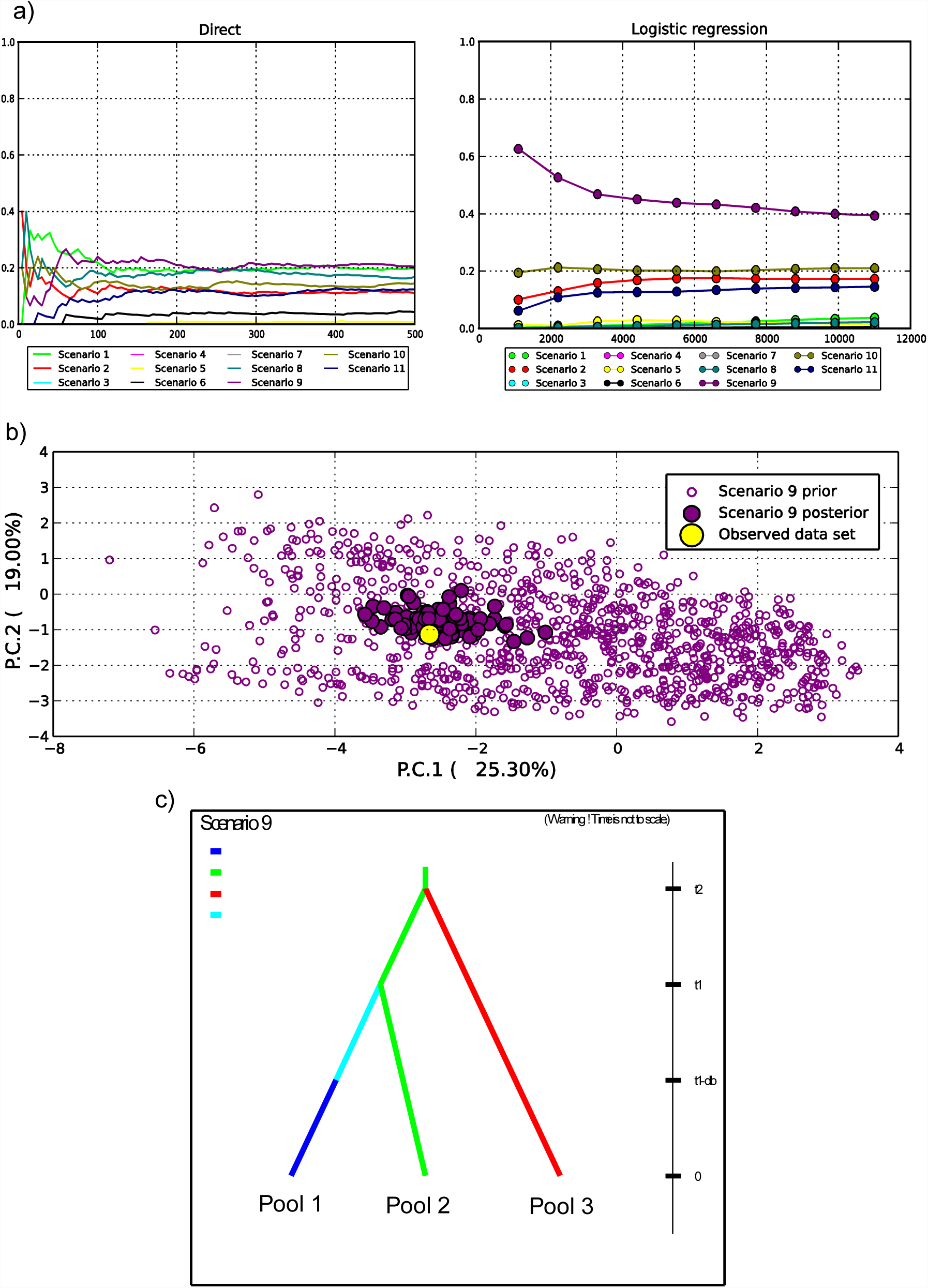
Broad scale DIYABC analysis (Stage 1) results. **a)** Direct approact (left) and Logistic regression (right) showing support for scenario 9. **b)** Model checking for scenario 9, showing that the observed data fall well within the cloud of datasets simulated from the posterior parameter distribution. **c)** Scenario 9 schematic.

**Supplementary Figure 9.**
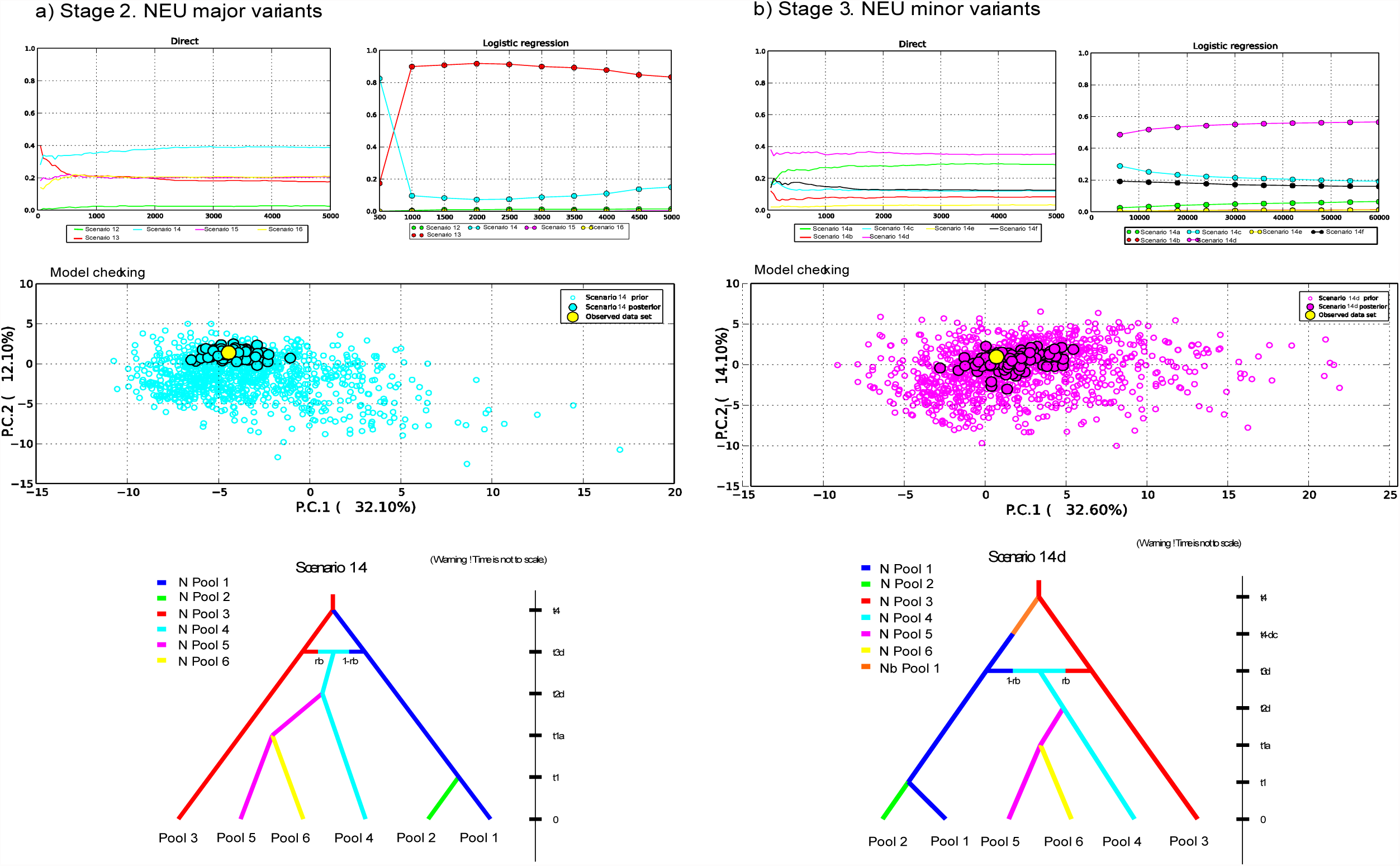
Fine scale DIYABC analysis in northern Europe‥ a) Stage 2 - major topological variants of scenarios. Direct approact (top left) and Logistic regression (top right) showing support for scenario 14 and 13 respectively. Model checking (Middle) for scenario 14 (bottom), showing that the observed data fall well within the cloud of datasets simulated from the posterior parameter distribution. Note the model checking placed the observed data outside of the cloud of posterior datasets for scenario 13. b) Stage 3 - Minor scenario variants of scenario 14 from stage 2. Direct approach (top left), logistic regression (top right) and model checking (middle) all support scenario 14d (bottom).

**Supplementary Figure 10.**
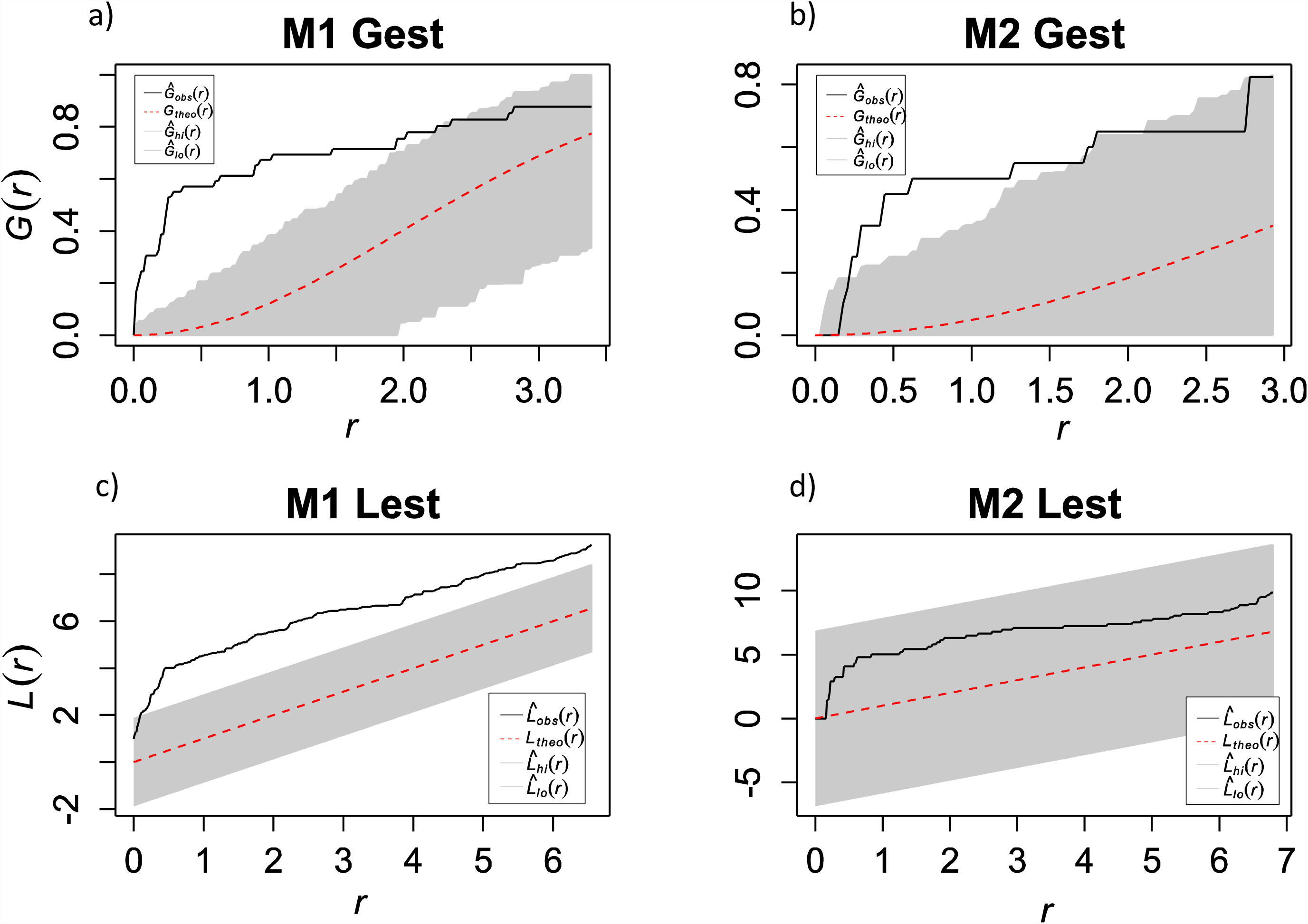
Comparison of spatial patterns of uniformity in geographic sampling regimes of the full M1 dataset locations (a, c) and the sampling location subset used in M2, M3, and RAD datasets (b,d). Estimates of G and L from true sampling locations are plotted using the black solid lines. Estimates of G and L from simulated locations based on random poisson distribution is represented by the red dashed line. Grey shaded areas are the 95% confidence intervals around the random estimates. Both the G and L function estimates show that there is more clustering of sampling locations in the M1 dataset than in the M2, M3 and RAD subsets.

